# Protective intravenous BCG vaccination induces enhanced immune signaling in the airways

**DOI:** 10.1101/2023.07.16.549208

**Authors:** Joshua M. Peters, Edward B. Irvine, Jacob M. Rosenberg, Marc H. Wadsworth, Travis K. Hughes, Matthew Sutton, Sarah K. Nyquist, Joshua D. Bromley, Rajib Mondal, Mario Roederer, Robert A. Seder, Patricia A. Darrah, Galit Alter, JoAnne L. Flynn, Alex K. Shalek, Sarah M. Fortune, Bryan D. Bryson

## Abstract

Intradermal (ID) Bacillus Calmette–Guérin (BCG) is the most widely administered vaccine in the world. However, ID-BCG fails to achieve the level of protection needed in adults to alter the course of the tuberculosis epidemic. Recent studies in non-human primates have demonstrated high levels of protection against *Mycobacterium tuberculosis* (*Mtb*) following intravenous (IV) administration of BCG. However, the protective immune features that emerge following IV BCG vaccination remain incompletely defined. Here we used single-cell RNA-sequencing (scRNAseq) to transcriptionally profile 157,114 unstimulated and purified protein derivative (PPD)-stimulated bronchoalveolar lavage (BAL) cells from 29 rhesus macaques immunized with BCG across routes of administration and doses to uncover cell composition-, gene expression-, and biological network-level signatures associated with IV BCG-mediated protection. Our analyses revealed that high-dose IV BCG drove an influx of polyfunctional T cells and macrophages into the airways. These macrophages exhibited a basal activation phenotype even in the absence of PPD-stimulation, defined in part by IFN and TNF-α signaling up to 6 months following BCG immunization. Furthermore, intercellular immune signaling pathways between key myeloid and T cell subsets were enhanced following PPD-stimulation in high-dose IV BCG-vaccinated macaques. High-dose IV BCG also engendered quantitatively and qualitatively stronger transcriptional responses to PPD-stimulation, with a robust Th1-Th17 transcriptional phenotype in T cells, and augmented transcriptional signatures of reactive oxygen species production, hypoxia, and IFN-γ response within alveolar macrophages. Collectively, this work supports that IV BCG immunization creates a unique cellular ecosystem in the airways, which primes and enables local myeloid cells to effectively clear *Mtb* upon challenge.

## INTRODUCTION

Bacillus-Calmette-Guerin (BCG) is administered to approximately 130 million children every year to protect against *Mycobacterium tuberculosis* (*Mtb*), the causative agent of tuberculosis (TB) (*1–3*). While intradermal Bacillus-Calmette-Guerin (BCG) is the only licensed TB vaccine, it inadequately protects against pulmonary infection in adults (*4*). Consequently, TB remains a leading driver of death worldwide (*5*). The development of an effective TB vaccine has been hampered by a poor understanding of protective immunity against *Mtb* (*6–8*). Intravenous (IV) administration of BCG was recently shown to be highly protective against *Mtb* in non-human primates (NHPs), making high-dose IV BCG immunization a powerful system to identify immune correlates and mechanisms of protection against *Mtb* infection (*9*, *10*).

Several immune features are associated with IV BCG vaccination (*8*, *9*, *11*, *12*). IV BCG immunized rhesus macaques exhibit higher antibody titers and a broad increase in CD4+ and CD8+ T cell immunity in the airways (*11*, *12*). Further, lung-localized T cells in IV BCG immunized macaques are enriched for a transcriptional signature of memory T cell functionality compared to macaques that received intradermal BCG or aerosol BCG vaccination (*9*). However, the phenotypic effect of IV BCG vaccination on the larger cellular ecosystem in the lungs and airways has not been evaluated. For instance, macrophages represent a central node in the immune cell network against *Mtb*, which can be modulated by other immune effectors such as T cells and antibodies to promote intracellular *Mtb* control (*13*, *14*). Despite evidence that the ability of macrophages to respond to and control *Mtb* infection is a function of their phenotypic state (*15*), the impact of IV BCG on airway myeloid cell phenotype and function in NHPs remains largely unknown.

In the present study, we leveraged single-cell transcriptional data from several cohorts of BCG vaccinated NHPs to develop an integrated view of how protective and non-protective vaccination impacts the airway immune cell environment. We hypothesized that profiling this unique resource of airway cells across multiple vaccine routes of administration and doses would reveal potential immune correlates of protection against *Mtb* and provide additional insights into mechanism(s) of IV BCG-induced protection. High-dose IV BCG induced a robust increase in T cells and recruited macrophages into the airways. Strikingly, high-dose IV BCG also drove a prolonged state of transcriptional activation in airway macrophages, characterized by robust IFN and TNF-α signaling that was detected up to 6 months following vaccination. Moreover, intercellular communication between select myeloid and T cell populations was significantly increased in the BAL of high-dose IV BCG immunized macaques upon stimulation with purified protein derivative (PPD), highlighting a broad increase in responsiveness to PPD-stimulation. Finally, transcriptional signatures of key antimicrobial processes including reactive oxygen species production, hypoxia, and IFN-γ response were selectively upregulated in alveolar macrophages upon PPD-stimulation in the high-dose IV BCG vaccination group. Taken together, this work suggests that IV BCG immunization establishes a milieu of highly coordinated immune cells in the lung that robustly activates local myeloid cells to counter *Mtb* infection.

## RESULTS

### An integrated census of bronchoalveolar lavage cells from BCG-vaccinated macaques

Bronchoalveolar lavage (BAL) cells from 29 rhesus macaques across multiple BCG vaccination cohorts were transcriptionally profiled using scRNAseq (Fig. 1A-B, Fig. S1A-D and Table S1). The first cohort consists of 15 total macaques immunized with low-dose intradermal (ID) BCG (BCG dose 5×10^5^ colony forming units (CFU); n = 3), high-dose ID BCG (BCG dose 5×10^7^ CFU; n = 3), high-dose aerosol BCG (BCG dose 5×10^7^ CFU; n = 3), or high-dose IV BCG (BCG dose 5×10^7^ CFU; n = 3), as well as naive, unvaccinated controls (n = 3) (*9*). BAL cells from these macaques were collected 13- and 25-weeks following vaccination and these animals were not challenged with Mtb. The second cohort contains 14 macaques vaccinated with IV BCG across 4 different doses (13 IV BCG, 1 naïve; doses: 1.37×10^5^ – 4.8×10^6^ CFU). BAL cells were collected from these macaques 14-15 weeks following vaccination. Ten macaques in the second cohort were later challenged with low-dose *Mtb* and bacterial burdens were measured 12 weeks post-infection.

**Fig. 1.**
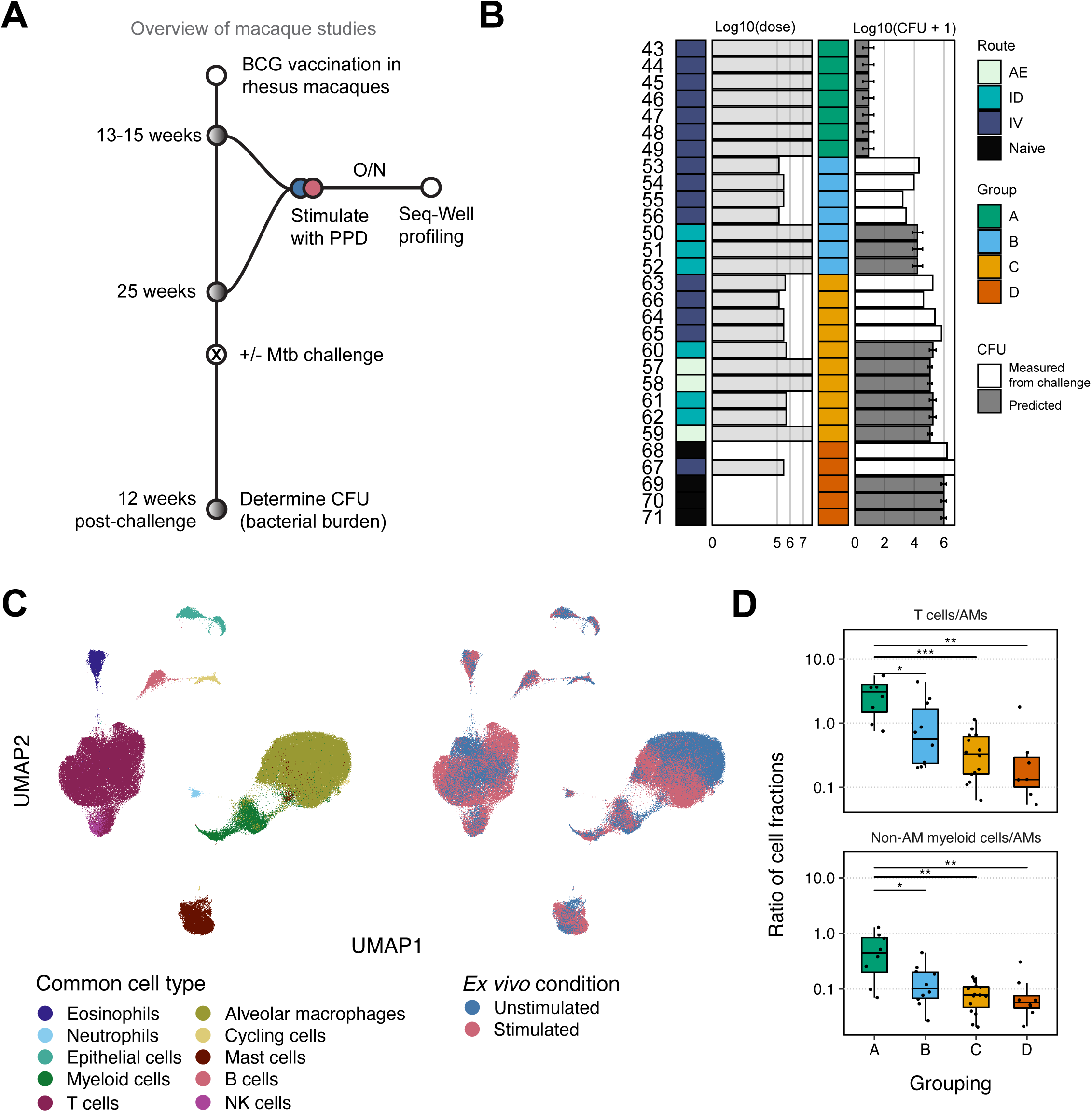
Integrated census of bronchoalveolar lavage cells displays vaccine-dependent shifts in immune cell populations. (**A**) Diagrammatic overview of study workflow, (**B**) Visual table of macaque samples describing the vaccine route (AE: aerosol, ID: intradermal, IV: intravenous, Naive: not vaccinated), log10(BCG vaccine dose), grouping (group A is estimated as most protected to group D as least protected), log10(CFU + 1) of macaques (estimated from cohort statistics for macaques without direct measurement). Error bars on estimated CFUs represent the average of historical CFUs ± 1 standard deviation. (**C**) UMAP embedding of all cells, colored by cell type and stimulation condition. (**D**) Ratio of cell fractions (as a fraction of total cells) between alveolar macrophages (AMs) and other immune cell types: T cells and myeloid cells. Each dot indicates a batch. Mann-Whitney *U* tests were performed between groups A/B, A/C, A/D, B/C, and C/D on the ratios of cell proportions per group of cell types. P values were adjusted using the Benjamini-Hochberg procedure. Adjusted P values are denoted by: *, p < 0.05; **, p < 0.01; ***, p < 0.001; ****, p < 0.0001. Only comparisons significant at p < 0.05 are shown.

BAL samples for each macaque at each timepoint were analyzed using Seq-Well (*16*, *17*), yielding 157,114 cells from 29 macaques after quality control and single cell data integration using Harmony (Fig. 1C, Table S2, Fig. S1E and S1F and Materials and Methods) (*18*). We identified cell states in the unstimulated (n = 74,641 cells) and PPD-stimulated (n = 82,473 cells) BAL cells together after harmonization using cluster markers and established lung cell type signatures (Fig. 1C, Fig. S1E and S1F, Fig. S2A, Table S4, and Materials and Methods) (*19*). Among these cells, we identified 53,979 T cells, 1,837 NK cells, 70,130 alveolar macrophages (AMs), and 8,815 non-alveolar macrophage myeloid cells (MCs), including monocytes, dendritic cells, and macrophages. We performed sub-clustering in the T/NK cells and AMs/MCs, given the large abundance of these cell types and known diversity within these cell lineages (Table S3). Cells were equally represented across batches, mitigating concern of batch-to-batch variation (Local Inverse Simpson’s Index, LISI = 8.70, Table S5, and Materials and Methods). Further, all cell types were captured in both unstimulated and PPD-stimulated samples (Fig. S2B).

To test the hypothesis that different BCG vaccination strategies drive unique cellular ecosystems in the lung, we first needed to group the macaques based on relevant biological variation. Not all the BCG-vaccinated macaques in this study were challenged with *Mtb*, so we were unable to simply group the macaques based on experimentally determined *Mtb* burden at a common timepoint. We therefore utilized both measured and historical average *Mtb* burdens to establish outcome groups. We classified each rhesus macaque into one of four groups, ordered A through D, based on their measured or historical average *Mtb* burden (CFU) at necropsy (Fig. 1B). We used the historical average *Mtb* burden determined by Darrah et al (Materials and Methods, Fig. 1B, Table S1 and Table S3) for macaques without necropsy *Mtb* burden data (*9*). Group A is comprised exclusively of macaques that received high-dose IV BCG, and thus is associated with the lowest *Mtb* burden (log10(μ_A CFU_) = 1.1). Group B includes four low-dose IV BCG macaques and three high-dose ID macaques (log10(μ_B CFU_) = 4.04). Group C has four low-dose IV BCG macaques, three low-dose ID macaques, and three high-dose aerosol macaques (log10(μ_c CFU_) = 5.16). Finally, Group D is comprised of four naïve macaques and one low-dose IV BCG macaque (log10(μ_D CFU_) = 6.1), and thus is associated with the highest *Mtb* burden. Comparisons across groups can be interpreted as a comparison of immune responses in the lung as an approximate function of historical and measured *Mtb* CFU burdens.

### Increased recruited macrophages in the BAL of high-dose IV BCG vaccinated macaques

Flow cytometry analysis previously showed that high-dose IV BCG vaccination drives a dramatic influx of T cells into the airways, resulting in a significant increase in the T cell:alveolar macrophage ratio in the BAL (*9*). Likewise, comparing cellular composition of the BAL across Groups A-D by scRNAseq revealed that ratio of T cells to alveolar macrophages was significantly higher in Group A compared to Groups B-D (Fig. 1D and Table S5). Unexpectedly, the ratio of non-alveolar macrophage myeloid cells to alveolar macrophages was also significantly higher in Group A than in the other groups, pointing to a compositional difference in the myeloid compartment associated with bacterial control (Fig. 1D and Table S5). These compositional differences were also significant when comparing high-dose IV BCG, low-dose IV BCG, and all other samples (Fig. S2C). Furthermore, these trends which discriminated Group A from Group D were consistent at 13 and 25 weeks (Fig. S2D).

Given that myeloid cells play a well-established role in *Mtb* containment (*13*, *14*), we next focused on identifying the myeloid cell clusters in unstimulated BAL cells as a function of group. Subclustering the AM/MC populations revealed six primary cell types, namely conventional type I dendritic cells (cDC1s), plasmacytoid DCs (pDCs), mregDCs, monocytes, alveolar macrophages, and recruited macrophages (Fig. 2A, Fig. S3A, and Table S3). To distinguish between recruited and resident macrophages, we used lineage-resolved signatures of recruited monocytes-macrophages in mice and humans (*20*) to annotate each group (Materials and Methods and Fig. S3B). Recruited macrophages lacked tissue-resident alveolar macrophage markers and expressed chemotactic markers such as *CXCL9/10/11*, as well as genes implicated in driving adaptive immune trafficking and activation (Table S3). Our compositional analysis revealed that unstimulated BAL samples in Group A had a significant positive enrichment of recruited macrophages and cDC1s, while alveolar macrophages were significantly negatively enriched (Fig. 2B and Table S4). These data suggest that beyond driving increased T cell numbers in the BAL, high-dose IV BCG vaccination also promotes an increase in macrophages into the airways, significantly shifting the local myeloid cell ecosystem.

**Fig 2.**
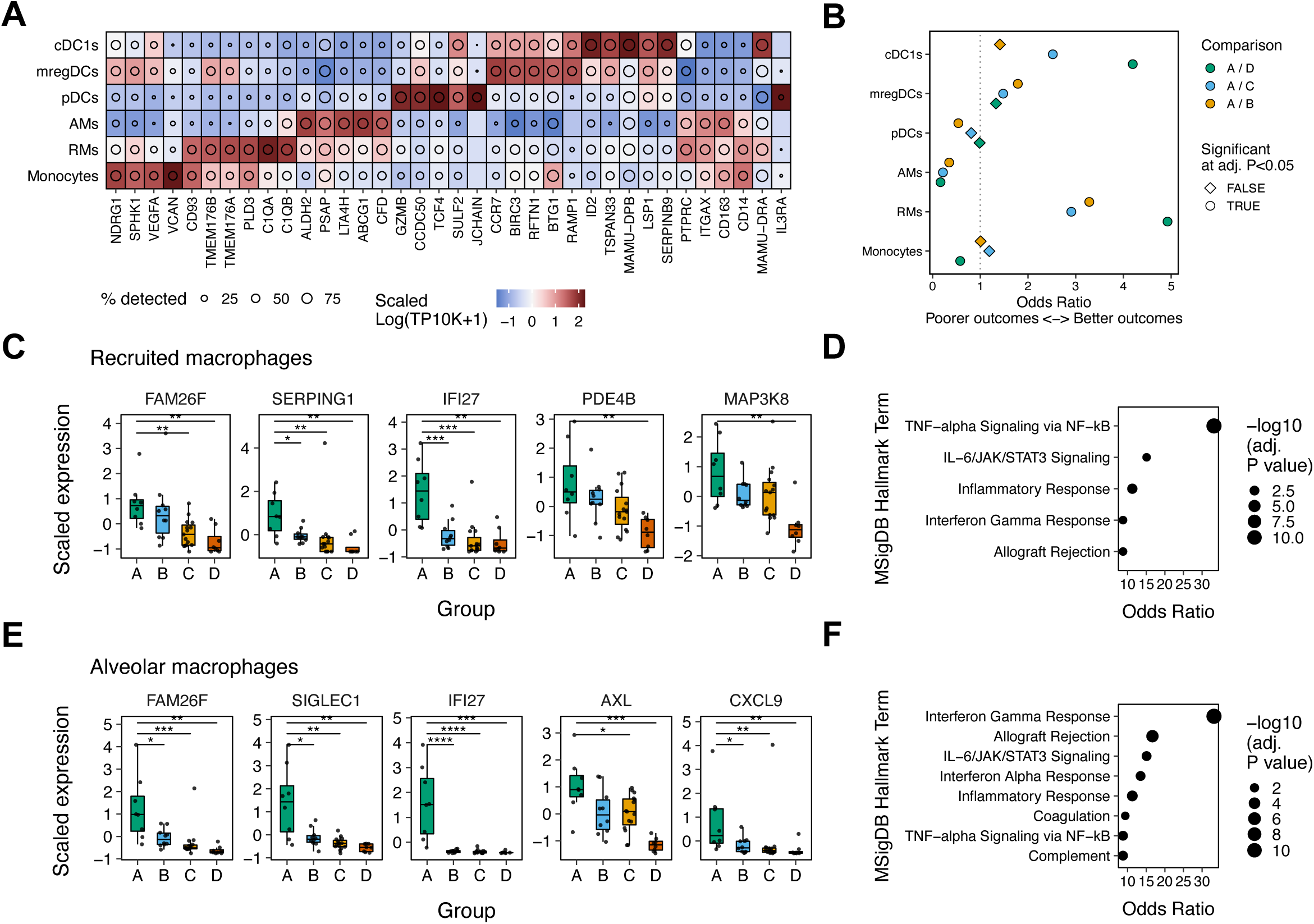
Recruitment and activation of myeloid cells in high-dose IV BCG macaques. (**A**) Scaled log-normalized expression of selected genes across annotated cell states. Circles within each tile of the heatmap represent the percentage of cells with the gene detected. The top 3 genes by AUROC for each cell type are selected, in addition to a panel of genes encoding protein targets typically used in flow cytometric panels (PTPRC to IL3RA). (**B**) Odds ratio for selected myeloid cell types comparing A/D, A/C, and A/B. Odds ratios are calculated from a binomial generalized linear model analyzing the number of cells for each cell type and sample and the total number of other cells within that sample as a function of cell type and quaternary group. (**C**) Scaled log-normalized expression (log10(TP10K +1)) values for selected genes within recruited macrophages in unstimulated samples across quaternary groups. Mann-Whitney *U* tests were performed between groups A/B, A/C, and A/D on the scaled expression values of each gene. (**D**) Enrichment of top 50 genes associated with quaternary groups based on Kendall’s τ using the MSigDB database and Enrichr. Odds ratio and -log10(adjusted P value) is plotted for each term significant at p < 0.05. **(E-F)** Same as (C) and (D) but for alveolar macrophages. P values were adjusted using the Benjamini-Hochberg procedure. Adjusted P values are denoted by: *, p < 0.05; **, p < 0.01; ***, p < 0.001; ****, p < 0.0001. Only comparisons significant at p < 0.05 are shown.

### High-dose IV BCG drives the robust activation of local macrophages

We next aimed to determine whether high-dose IV BCG primed a difference in transcriptional state in the lung-localized macrophages that may contribute to altered immune ecosystems following vaccination. In recruited macrophages, Mann-Whitney analysis identified activation-associated genes, such as *FAM26F* and *IFI27*, to be expressed at significantly higher levels in Group A in comparison to Groups C and D in unstimulated cells (Fig. 2C, Table S5, and Materials and Methods). We next performed gene set enrichment using the 50 genes displaying the strongest correlation with group in unstimulated recruited macrophages, based on the Kendall rank correlation coefficient (Kendall’s τ). Kendall’s τ was used due to the ordinal nature of groups A though D. This analysis found TNF-α, IL-6, and IFN-γ signaling pathways, underscored by *IL1B*, *NLRP3*, *TLR2*, and *PDE4B* expression, to be significantly positively enriched (Fig. 2D), highlighting the pro-inflammatory transcriptional profile of recruited macrophages particularly in the airways of Group A macaques.

Previous reports suggest that alveolar macrophages are highly permissive for *Mtb* since their activation is tightly regulated to limit undesired inflammatory responses (*21*). We found expression of immune activation genes in unstimulated AMs that also tracked with group (Fig. S3C-D and Table S5). For instance, genes such as *FAM26F* and *IFI27* which exhibited increased expression in the recruited macrophages of Group A (Fig. 2C), were also significantly more highly expressed in Group A alveolar macrophages compared to alveolar macrophages in the other groups (Fig. 2E, Table S5). Moreover, gene set enrichment using the 50 genes correlated with protection in unstimulated alveolar macrophages (Kendall’s τ; Materials and Methods) found a significant positive enrichment of inflammatory pathways including IL-6 signaling, the type I and type II IFN response, and the TNF-α response (Fig. 2D), driven by genes such as *CD86, CXCL9, STAT1,* and *TLR2* (Fig. 2F, Table S5). Conversely, *PTGDS*, which encodes a synthase responsible for PGD2 production and inflammation dampening, was strongly correlated with Group D (*22*). Taken together, these data point to a marked compositional shift in the local macrophage compartment up to 6 months following high-dose IV BCG vaccination via two mechanisms: (i) the influx of inflammatory recruited macrophages into the airways, and (ii) the phenotypic shift of resident alveolar macrophages to a protracted state of activation.

### High-dose IV BCG enhances myeloid-T cell communication

The airway of IV BCG immunized macaques represents a unique multicellular environment characterized by a network of immune cell populations whose collective activity likely influences myeloid cell action and TB disease trajectory (*23*, *24*). Thus, to infer interaction between distinct immune cell populations, we performed cell-cell interaction analyses using CellChat (Materials and Methods). CellChat infers interaction strengths between cell subsets using ligand-receptor expression and cellular composition information (*25*).

The total interaction strength, representing incoming and outgoing ligand-receptor interactions, revealed cells in the AM and MC compartments as dominant signal receiver and sender populations in unstimulated cells (Fig. 3A, left). While these cellular subsets remained dominant signal receiver and sender populations upon PPD-stimulation, polyfunctional T cells, helper CD4+ T cells, cDC1s, and mregDCs also emerged as potent signal sender and/or receiver populations following stimulation (Fig. 3A, right).

**Fig. 3.**
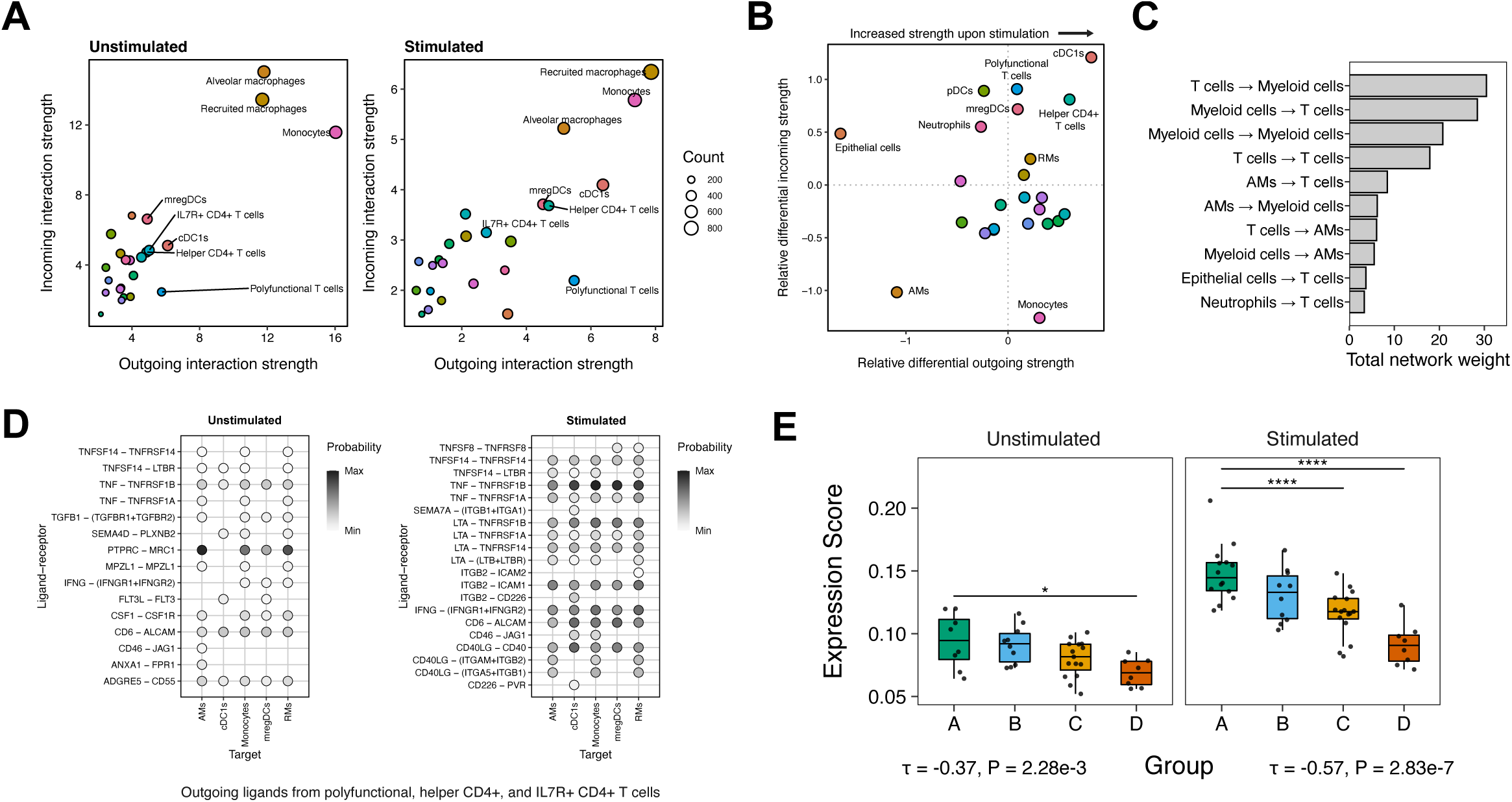
PPD-responsive T cell-myeloid interactions. (**A**) Absolute (left) and relative (right) changes in outgoing and incoming interaction strength, a measure of ligands’, receptors’, and associated molecules’ expression and cell proportions, for all cell states between unstimulated and PPD-stimulated samples. Positive values signify increases in interaction strength upon *ex vivo* stimulation. (**B**) Relative interaction strengths for all cell states between unstimulated and PPD-stimulated samples. **(C)** Total network weight within the cell-cell communication network between cell types grouped by their broader lineage. (**D**) Interaction strength of ligand-receptor interactions between polyfunctional, activated, and helper CD4+ T cells and selected myeloid cells for unstimulated cells (left) and PPD-stimulated cells (right). (**E**) Expression score of top 20 ligand-receptors pairs across groups. Mann-Whitney *U* tests were performed between groups A/B, A/C, and A/D on the total expression score. P values were adjusted using the Benjamini-Hochberg procedure. Adjusted P values are shown directly.

Given that polyfunctional and helper CD4+ T cells exhibited a robust increase in intercellular communication following PPD-stimulation (Fig. 3B), we next interrogated the likely signaling partners of these key populations. T cell-myeloid interactions dominated intercellular signaling overall (Fig. 3C). In unstimulated cells, these subsets primarily interacted through CD6-ALCAM, CSF1-CSF1R, TNF-TNFRSF1B, as well as FLT3L-FTL3 and IFNG-IFNGR1/2 (Fig. 3D, left).

Notably, CD6-ALCAM and FLT3L-FLT3 interactions are implicated in driving T cell effector function and dendritic cell activity (*26–29*). Upon PPD-stimulation, we see an increase in other ligand-receptor interactions involved in T cell-induced macrophage activation including CD40LG-CD40 and LTA-TNFRSF1B (Fig. 3D, right), as well as increased TNF and IFNG interactions.

We next investigated how ligand-receptor interactions varied across outcome groups. For each ligand-receptor pair within the top signal sending and receiving populations, we calculated the correlation of their expression with outcome group (Table S6). The top 15 ligand-receptor interactions displaying the strongest correlation with outcome group comprised 24 genes, including *CD40*, *CD80*, *CD28*, *LTA*, *CCL20*, *TNF*, *IFNG*, and their cognate partners. Scoring samples based on the expression of these key ligand-receptor pairs revealed a significantly higher expression score in Group A macaques compared to those in Group D in unstimulated cells (Fig. 3E). In PPD-stimulated samples, expression scores in Group A macaques were even higher (Fig. 3E), pointing to a particularly robust increase in T cell-myeloid signaling through ligands such as TNF, IFNG, and CD40L upon PPD-stimulation in high-dose IV BCG vaccinated macaques. These trends were consistent at 13- and 25-weeks following vaccination (Fig. S4). Together, these findings identify myeloid cells as key signal sender and receiver populations in the BAL of BCG immunized macaques, and further suggest that high-dose IV BCG vaccination primes an airway environment that promotes increased communication of myeloid cells with polyfunctional and helper CD4+ T cells upon antigenic stimulation.

### Myeloid-interacting T cells display increased T1-T17 polarization following high-dose IV BCG vaccination

Given the strong predicted communication between AM/MCs and T cells, we next examined the compositional and transcriptional differences in the airway T/NK cell compartment with increased granularity. Subclustering of the T and NK cell populations identified 13 distinct cell subsets (Fig. 4A-B and Materials and Methods), including *IFNG* and *TNF*-expressing polyfunctional T cells (n = 1,460), and *IRF4* and *CD28*-expressing helper *CD4*+ T cells (n = 6,540). Several transcription factors implicated in T cell activation and polarization were defining markers of cell states, including *IRF4, RBPJ, BATF, NR4A3*, and *EGR2* (*30–37*). To determine which T/NK cell subsets exhibited the strongest associations with group, we compared the proportional abundance of each subset across outcome groups (Fig. S5A-C, Table S4, and Materials and Methods). The proportional abundance of nearly all T cell subsets significantly correlated with group, with Tregs being the only subset with no significant association with group (Kendall’s τ = 0.03) (Fig. S5C). Of note, in both unstimulated and PPD-stimulated BAL cells, polyfunctional T cells (mean Kendall’s τ = -0.63), helper CD4+ T cells (mean Kendall’s τ = -0.57), and IL7R+ CD4+ T cells (mean Kendall’s τ = -0.48) – each identified previously as primary myeloid interaction partners (Fig. 3A) – displayed particularly strong correlations with group (Fig. 4C).

**Fig 4.**
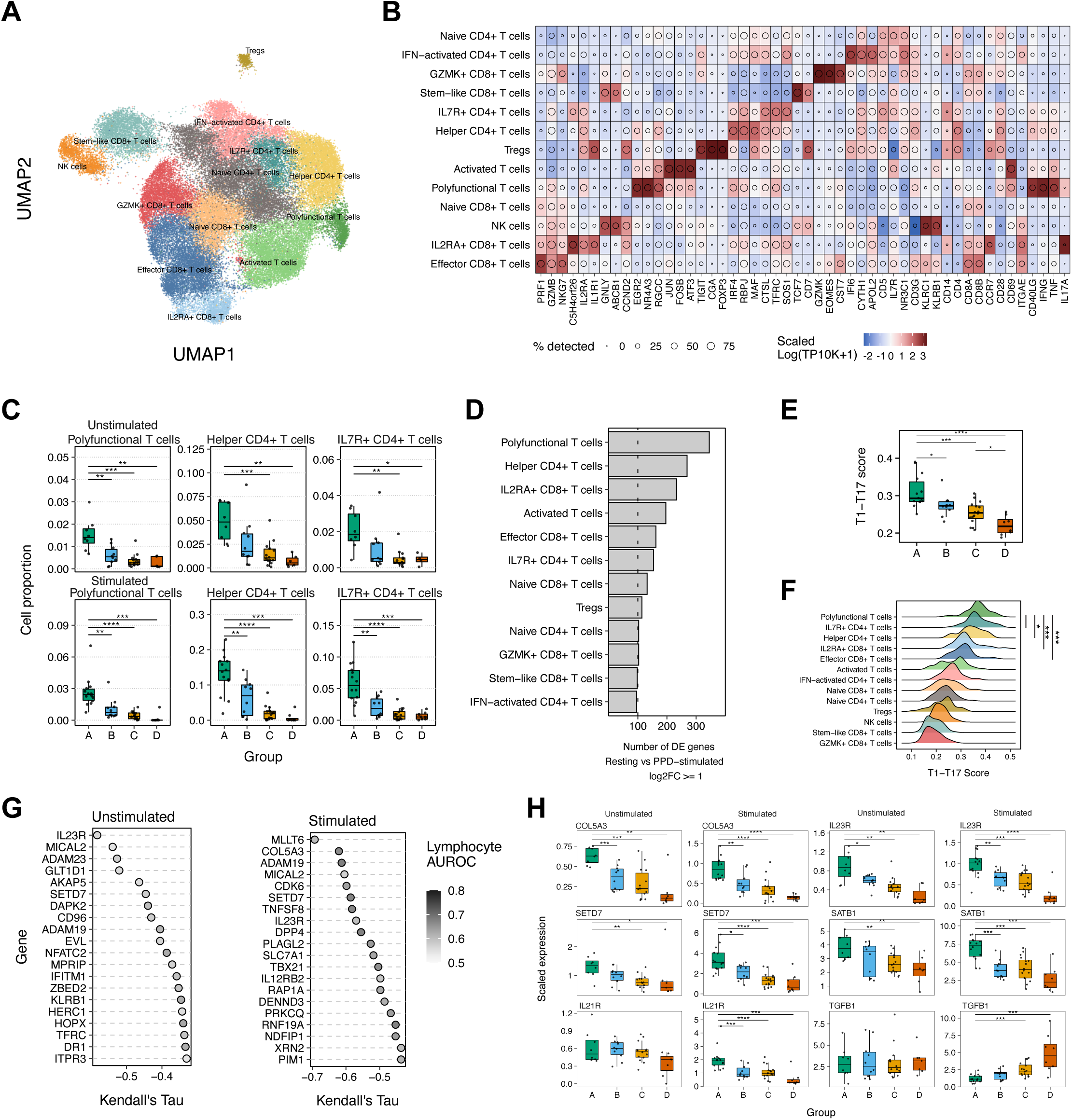
Polyfunctional T cell populations are most associated with protection. (**A**) UMAP embedding of T and NK cells, colored by subclustered cell state annotations. (**B**) Scaled log-normalized expression of selected genes across annotated cell states. Circles within each tile of the heatmap represent the percentage of cells with the gene detected. The top 3 genes by AUROC for each cell state are selected, in addition to a panel of genes encoding protein targets typically used in flow cytometric panels (CD3G to IL17A). (**C**) Cell proportions of total cells for selected cell states in both unstimulated and PPD-stimulated samples across quaternary groups. Mann-Whitney *U* tests were performed between groups A/B, A/C, and A/D on the cell proportions of total cells for each cell state. P values were adjusted using the Benjamini-Hochberg procedure. (**D**) Number of differentially expressed genes between unstimulated and PPD-stimulated samples within each T cell subset. Only genes with log2(fold change) >= 1 included. (**E**) T1-T17 scores across T and NK cells within stimulated samples. The T1-T17 signature genes are derived from Gideon *et al.* 2021. Mann-Whitney *U* tests were performed between A/B, A/C, A/D, B/C, and C/D. **(F)** T1-T17 scores across T and NK cells within all samples. Cell states are ordered by mean score from high to low. Mann-Whitney *U* tests were performed between the first and subsequent four cell states. **(G)** Gene Kendall’s Tau correlations with Group. Points are colored by the AUROC of each gene describing the specificity of the gene to a cell state within the T and NK cell population. **(H)** Log-normalized expression values for selected genes within all T and NK cells across quaternary groups split by stimulation state. P values were adjusted using the Benjamini-Hochberg procedure. Adjusted P values are denoted by: *, p < 0.05; **, p < 0.01; ***, p < 0.001; ****, p < 0.0001. Only comparisons significant at p < 0.05 are shown.

To more precisely characterize the functionality and responsiveness of the different BAL T cell subpopulations, we asked which T cells were most transcriptionally-responsive to PPD by analyzing the number of differentially-expressed genes (DEGs) upregulated after PPD stimulation per subset. Polyfunctional and helper CD4+ T cells were most responsive to PPD-stimulation based on the number of differentially-expressed genes, followed by IL2RA+ CD8+ T cells, activated T cells, and effector CD8+ T cells (Fig. 4D). We next contextualized these T cell populations using recent and biologically relevant single-cell RNA-sequencing-derived signatures. Specifically, following *Mtb* infection and the onset of adaptive immunity, a “T1-T17” population was found to be highly enriched in *Mtb* granulomas that promote bacterial clearance (*13*). To determine whether a similar transcriptional signature was engendered in the setting of high-dose IV BCG vaccination, we scored T and NK cells on the top 50 genes defining this T1-T17 population. Indeed, the T1-T17 transcriptional signature was significantly enriched in Group A macaques across all T/NK cell populations (Fig. 4E). This broad shift across diverse T/NK cell subtypes towards a T1-T17 phenotype suggests that in this context the T1-T17 phenotype may represent a polarization state driven by high-dose IV BCG vaccination, rather than a specific subset of T cells. Here, the key myeloid-interacting T cell subsets – polyfunctional T cells, helper CD4+ T cells, and IL7R+ CD4+ T cells – exhibited the strongest T1-T17 score (Fig. 4F). To further define T cell phenotypic state, we scored T and NK cells using a signature of lung residency containing 10 genes derived from the human lung atlas (Materials and Methods) (*19*). We observed significantly higher expression of lung residency-defining genes within the polyfunctional and helper CD4+ T cell subsets and lower expression in naïve and stem-like subsets (Fig. S6A).

We next sought to uncover gene-level correlates with group (Table S7) (Materials and Methods). In unstimulated T cells, *IL23R*, implicated in Th1 and Th17 proliferation, and *CD96*, a co-stimulatory and immunoregulatory receptor, were among the top genes associated with group (Fig. 4G, left) (*38–41*). Within PPD-stimulated T cells, a range of functionally-relevant genes such as *COL5A3*, a Th17 marker (*42*), *TNFSF8* (CD30LG, CD153), indicative of IL-17 secretion (*43*), and *DPP4*, associated with MAIT cells (*44*), significantly associated with group (Fig. 4G, right). Furthermore, the comparison of gene expression profiles across groups revealed qualitatively distinct expression patterns between unstimulated and PPD-stimulated samples. For instance, *COL5A3, IL23R, SETD7,* and *SATB1* represented examples of genes with increased expression in Group A across unstimulated and PPD-stimulated samples. Yet genes such as *IL21R*, implicated in CD8+ T cell priming and maintenance under chronic stimuli (*45*, *46*), only showed significantly higher expression in Group A upon PPD-stimulation. Conversely, *TGFB1*, previously implicated in *Mtb* immunosuppression (*47*), showed little specificity to any particular T cell subset in unstimulated samples. However, upon PPD stimulation, *TGFB1* was significantly more expressed in Group D (Fig. 4H). Collectively, these data indicate that beyond broad increases in T cell abundance, specific transcriptional programs in T/NK cells may regulate protective (T1-T17) or misdirected (*TGFB1*) immunity to *Mtb* infection respectively.

### Select super-induced genes upon PPD-stimulation track with outcome group

Distinct expression patterns between unstimulated and PPD-stimulated T cells inspired the hypothesis that quantitative and/or qualitative differences in cellular responsiveness to PPD-stimulation, an *ex vivo* proxy for *Mtb* challenge, may associate with outcome group. To investigate this possibility, we leveraged the paired unstimulated and PPD-stimulated samples from the BAL of each macaque to query the relationship between cellular responsiveness to PPD-stimulation and group (Fig. 5A).

**Fig 5.**
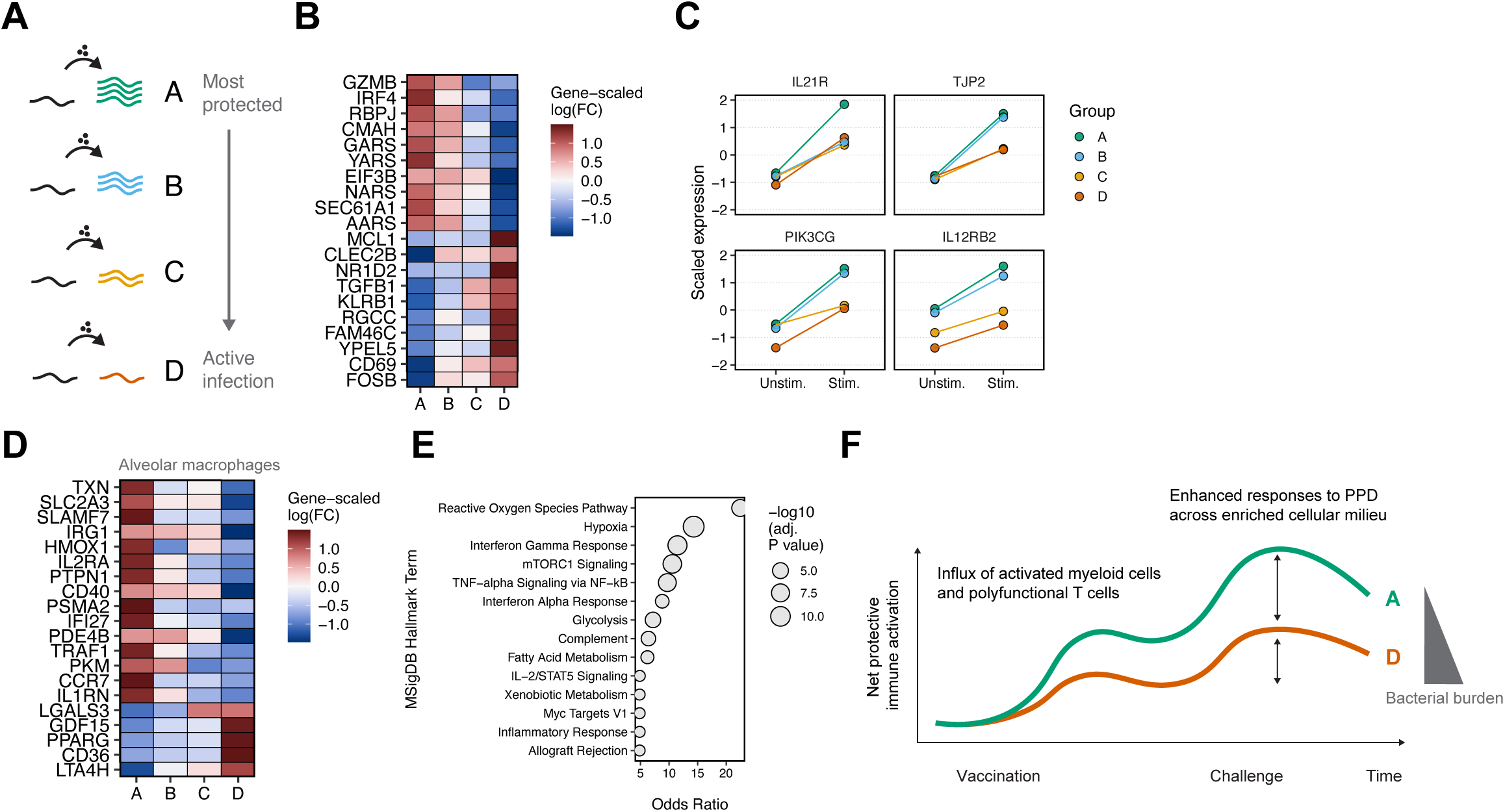
Responses to secondary antigen challenges are associated with protection. Alveolar T cell responses in (**B-C**) and macrophages responses in (**D-E**). (**A**) Analytical schematic depicting the identification of genes which are variably induced upon PPD stimulation based on group. **(B)** Heatmap of genes induced by PPD stimulation across quaternary groups. Color represents the gene-scaled log(fold change) of each gene in each group. **(C)** Scaled average expression per gene between unstimulated and PPD-stimulated samples across groups. **(D)** Heatmap of genes induced by PPD stimulation across quaternary groups. Color represents the gene-scaled log(fold change) of each gene in each group. **(E)** Enrichment of genes whose expression is induced by PPD in group-dependent manner using the MSigDB database and Enrichr. Odds ratio and -log10(adjusted P value) is plotted for each term significant at p < 0.05. (**F**) Schematic describing proposed model of lung microenvironment priming in Group A and Group D.

We constructed a PPD-response score using the top 50 genes upregulated following PPD-stimulation in polyfunctional, helper CD4+, and IL7R+ CD4+ T cells (Fig. S7A). These cell subsets were used to define the response signature as they displayed a strong and significant association with protective immunity, and therefore may represent a preferred response to antigenic stimulation. We then scored T cells across all subsets using this response signature, and then compared the fold change induction of this signature between unstimulated and PPD-stimulated samples. We observed a negative correlation across groups (mean Kendall’s τ = -0.39), with significantly stronger induction of PPD-upregulated genes in Group A as compared to Group D (Fig. S7B). This effect was seen across all T cell subsets except Tregs (Fig. S7C), indicating that the magnitude of the T cell transcriptional response to PPD-stimulation associated with group.

While the magnitude of the T cell response induced by antigen stimulation may represent a determinant of vaccine-induced protection, we reasoned that the type of response mounted may represent another critical feature. Hence, we next aimed to identify the genes that showed differential responses to PPD-stimulation across different outcome groups. We calculated the log(fold change) (logFC) in gene expression between unstimulated and PPD-stimulated T cells within each group (Table S8 and Materials and Methods) and identified genes which showed stronger fold changes in Group A. Transcriptional regulators such as *IRF4*, *RBPJ*, and *EIF3B* were selectively enriched upon PPD-stimulation in Group A (Fig. 5B) (*32*, *48*, *49*). Inversely, *TGFB1* and *KLRB1* were enriched in Group D, showing that PPD-stimulation induces disparate transcriptional responses in genes across outcome groups that mark or drive distinct T cell phenotypes. We further refined this question to identify genes whose expression in Group A (unstimulated and stimulated) was highest and had the highest fold change between conditions. These genes would be most elevated by high-dose IV BCG, and most responsive during a secondary challenge such as PPD-stimulation. We identified several genes matching these criteria: *IL21R, TJP2, PIK3CG,* and *IL12RB2*, among others (Fig. 5C). Critically, these genes have been implicated in local T cell persistence and function (*45*, *46*, *50–55*). These data describe core T cell processes mobilized upon antigenic stimulation in high-dose IV BCG vaccinated macaques.

Given the striking increase in T cell-myeloid communication upon PPD-stimulation identified in the ligand-receptor interaction analysis (Fig. 3A), we next sought to establish whether changes in unstimulated alveolar macrophages altered their response to secondary BAL stimulation with PPD *ex-vivo.* We generated a PPD-response signature by performing differential expression between paired unstimulated and PPD-stimulated bulk BAL samples (Materials and Methods). The signature included *STAT1* and *STAT1*-associated genes such as *CXCL9/10/11*, *GBP2*, and *CD274* (Fig. S3E). We found that the PPD-response signature score was elevated in unstimulated alveolar macrophages in Group A, indicating a state of baseline activation in the alveolar macrophage compartment of high-dose IV BCG immunized macaques that resembles activation induced by PPD-stimulation (Fig. S3F-G). We lastly sought to identify the genes that showed differential responses to PPD-stimulation across different outcome groups in alveolar macrophages. S*LAMF7*, a marker of macrophage super-activation (*56*, *57*), *SLC2A3*, a glucose transporter, *IL1RN*, an antagonist of IL-1 signaling (*58*), and *IRG1*, implicated in acute *Mtb* infection resolution (*59*), represented examples of genes selectively upregulated upon PPD-stimulation in Group A (Fig. 5D). Pathway analysis of these enriched genes indicated that AMs from high-dose IV BCG vaccinated macaques selectively and significantly mobilized biological processes that generate a hostile intracellular environment for the bacterium, such as reactive oxygen species, hypoxia, and the IFN-γ response following PPD-stimulation (Fig. 5E) (*7*). Together, these results suggest that high-dose IV BCG immunization primes a unique cellular environment in the lungs, defined by a heightened functional responsiveness to *Mtb* challenge that likely arms local immune cells to counter *Mtb* infection (Fig. 5F).

## DISCUSSION

In this study, we exploited single-cell transcriptional data from several cohorts of BCG vaccinated NHPs to discover how different TB vaccination paradigms which vary in their protective efficacy alter immune ecosystems in the lung. We found that high-dose IV BCG vaccination remodeled the airway myeloid compartment, triggering a sustained compositional shift towards inflammatory recruited macrophages and activated alveolar macrophages that serve as master communicators of intercellular signaling in the lung. Upon PPD-stimulation, intercellular communication between BAL macrophages and T cells was increased in high-dose IV BCG immunized macaques (Group A). This group also showed quantitatively and qualitatively enhanced transcriptional responses to PPD-stimulation, with a distinct Th1-Th17 transcriptional phenotype in T cells and augmented transcriptional signatures of reactive oxygen species production, hypoxia, and IFN-γ responses within the alveolar macrophage compartment, collectively suggesting that IV BCG primes a unique cellular ecosystem in the lung, which enables myeloid cells to effectively combat *Mtb* upon challenge.

In mice and humans, BCG vaccination results in the innate training of myeloid cells, characterized by a heightened responsiveness to stimuli and epigenetic remodeling (*60–65*). In light of this, our observation that alveolar macrophages displayed an enhanced transcriptional response upon stimulation with PPD *ex-vivo* was intriguing. Still, our data suggest that that myeloid cell priming may be an apt description of the impact of high-dose IV BCG vaccination on local macrophage cell state (*66*). While immune activation returns to a baseline level following initial stimulation in innate training, during innate immune priming, the initial stimuli trigger a sustained transcriptional activation state such that secondary challenge engenders an even more potent response (*66*). Consistent with this model, our data indicate that 6 months following IV BCG immunization, airway macrophages of high-dose IV BCG vaccinated macaques display a protracted activation phenotype through IFN and TNF-associated transcriptional programs that resembles activation by PPD-stimulation, and further that their heightened activation state results in an enhanced transcriptional response upon stimulation with PPD *ex-vivo*. Measuring either innate training and innate immune priming in this system of stimulating BAL with PPD is challenging, given that cell type specific transcriptional responses to PPD-stimulation likely represent an integration of cell-intrinsic, autocrine, and paracrine effects. Importantly, given that we observed an increase in activated T cells in the airway, it is essential that we recognized and consider the contribution of T cells to the altered myeloid cell states in addition to any classical innate immune training which may also be occurring. Further, we lack adequate temporal resolution in this experimental set-up. Nevertheless, this study motivates future work investigating local myeloid cell priming in the context of *Mtb* infection and vaccination.

Alveolar macrophages of high-dose IV BCG vaccinated macaques displayed increased induction of reactive oxygen species, hypoxia, and IFN-γ response transcriptional programs following PPD-stimulation, highlighting the diverse intracellular and intercellular immune pressure triggered by antigenic stimulation in the setting of high-dose IV BCG vaccination. Similarly, the magnitude of the transcriptional response to PPD-stimulation correlated with outcome group across nearly all T cell subsets, and intercellular communication between myeloid cells and T cells was enriched in high-dose IV BCG immunized macaques following PPD-stimulation. These observations reveal a heightened phenotypic responsiveness to PPD-stimulation in the airway immune cells of macaques receiving high-dose IV BCG vaccination, and further suggest that IV BCG primes a local biological network of immune cells, poised to efficiently respond to and mobilize anti-*Mtb* immune function upon challenge. We note that while macrophage markers were included in the previous flow cytometry characterization of BAL cells in IV BCG, scRNAseq analysis in this study enabled the discovery of novel signatures and markers that define differences in macrophage state that should be considered for inclusion in future flow panels.

While the frequency of nearly all T cell populations were associated with outcome group, the strongest, most robust association was observed in polyfunctional and helper CD4+ T cells – both enriched for tissue-resident gene programs. These T cell subsets also displayed the strongest ligand-receptor interactions with the recruited myeloid cell populations. We identified a range of transcription factors (TFs) co-expressed within these T cell subsets, including: *EGR2, NR4A3, IRF4,* and *RPBJ*, in addition to T-bet (*TBX21*) *BATF*, and *RORC*. Critically, *IRF4* and *BATF*, implicated in driving effector function and countering exhaustion (*32*, *67*, *68*), strongly defined protective states. These polyfunctional T cells included both CD4+ and CD8+ T cells, indicating a consistent transcriptional signal of T cell polyfunctionality across CD4 and CD8+ T cells that are associated with outcome group. Like recent work that identified the frequency of CD4 memory T cells producing TNF with IFN-γ, and TNF with IL-17, as key predictors of protection (*12*), this work found a Th1-Th17 polarization of T cells induced by protective vaccination. Furthermore, outcome group-associated T cells in the BAL from our study, such as the polyfunctional and helper CD4+ T cell subsets, dominantly expressed a “T1-T17’’ transcriptional signature previously found to be stronger in T cells present in *Mtb* granulomas with low bacterial burdens (*13*), highlighting a convergence of T1-T17 transcriptional programs that associate with beneficial immune responses pre- and post-*Mtb* infection. Consistent with this observation, T cell expression of *IL23R*, *IL21R*, *PIK3CG, TJP2*, among others, before and after stimulation were highly associated with high-dose IV BCG vaccination in these analyses. These genes have been consistently implicated in regulating optimal T cell differentiation and activation towards long-lived, effector subsets (*40*, *41*, *45*, *46*, *50–52*, *69*). Overall, IV BCG biases the T cell compartment towards durable, primed polyfunctional states that mount stronger responses and intercellular communication after antigenic challenge. These data provide novel resolution into the molecular features underpinning these states and point towards a combination of transcription factors and receptors that can be exploited for future studies.

Our work underscores the complex logic of sterilizing immunity, which may require both the presence of some key immunoregulatory molecules and the absence of others. For instance, these data highlight a potential anti-protective effect of TGFB1 expression by T cells, as *TGFB1* was selectively augmented upon PPD-stimulation in macaques unable to control *Mtb* infection (Group D). This pattern aligns with recent work demonstrating that TGFβ impedes T cell expansion and function in TB granulomas (*47*). Expression of *GDF15*, a member of the TGF family secreted by macrophages and associated with tissue tolerance, iron availability, and immunosuppression (*70–72*), was also increased in Group D. These observations offer two distinct examples of objectives in immune correlates analyses: understanding the immune determinants of failure, and understanding multicellular profiles to capture immune system dynamics more holistically (*73–75*).

Limitations of the study include that *Mtb* burden data was not available for every macaque in our analysis, and instead we leveraged historical average *Mtb* CFU burdens for vaccination paradigms where we did not have paired burden data (n = 42). For instance, we associated the high-dose IV BCG immunized macaques comprising Group A as protected given the near-sterilizing protection observed in this setting historically (*9*). However, the incomplete experimental outcome data may complicate this interpretation. Furthermore, data were combined from multiple independent BCG vaccination studies in this analysis which could introduce batch related artifacts, despite efforts to mitigate this through Harmony-based batch correction (*18*).

The strengths of the study lie in our unbiased and comprehensive exploration of immune signatures of high-dose IV BCG vaccination – the current gold standard for protective immunity against *Mtb*. More generally, these data provide a large reference of immune cell profiles in the lung after administration of a systemic immunogenic agent. Our analysis of genome-wide transcriptional dynamics in a pairwise manner between unstimulated and antigen stimulated samples further offers a novel approach to identify gene programs modulated by antigenic challenge that correlate with a spectrum of infection outcomes.

Ultimately, this study contributes to increasing evidence that high-dose IV BCG vaccination fundamentally remodels the lung ecosystem of immunized macaques (*9*, *11*, *76*), priming the multicellular coordination likely required for *Mtb* control (*13*, *76–78*). Future work should investigate the temporal and expression dynamics of the key biologic networks identified herein immediately post-BCG vaccination, pre-*Mtb* challenge, and post-*Mtb* challenge to elucidate their collective role in creating a primed lung milieu capable of driving sterilizing protection against *Mtb* to catalyze the development of novel vaccination strategies to combat tuberculosis.

## MATERIALS AND METHODS

### Study Design

Indian-origin rhesus macaques (Macaca mulatta) used in these studies are outlined in Supplementary Table 1. All experimentation complied with ethical regulations at the respective institutions (Animal Care and Use Committees of the Vaccine Research Center, NIAID, NIH). Macaques were housed and cared for in accordance with local, state, federal, and institute policies in facilities accredited by the American Association for Accreditation of Laboratory Animal Care (AAALAC), under standards established in the Animal Welfare Act and the Guide for the Care and Use of Laboratory Animals. Macaques were monitored for physical health, food consumption, body weight, temperature, complete blood counts, and serum chemistries. All infections were performed at the University of Pittsburgh where animals were housed in a biosafety level 3 facility.

Samples were classified into four groups, A through D, based on measured CFU at the time of necropsy (week 36), or based on historical average CFU for a given vaccine strategy for macaques that were not challenged with *Mtb* (*9*). Specifically, historical data of vaccination and challenge studies were utilized to estimate the average *Mtb* CFU outcome for each vaccination strategy (n = 42). This generated four groups based on these vaccination strategies from cohort 1: high-dose IV (log_10_(BCG dose) > 6.5, μ_log10(*Mtb* CFU)_ = 1.1) as group A, high-dose ID (μ_log10(*Mtb* CFU)_ = 4.201) as group B, low-dose ID (μ_log10(*Mtb* CFU)_ = 5.236), aerosol (μ_log10(*Mtb* CFU)_ = 5.040) as group C, and unvaccinated macaques (μ_log10(*Mtb* CFU)_ = 5.975) as group D. High-dose IV macaques (n=4) were classified into group A, unvaccinated and macaques with log_10_CFU > 5.92 were classified into group D. The remaining 8 macaques were split in group B and C based on the 50^th^ percentile log_10_CFU = 4.31. Overall, these cutoffs align with the bottom 10% (log_10_CFU = 0.15), bottom 40% (log_10_CFU = 4.6), and bottom 90% (log_10_CFU = 6.16) of *Mtb* CFU measurements historically (n = 42). These groups resulted in average *Mtb* CFU for each group: μ_A_=1.1, μ_B_=4.04, μ_C_=5.16, μ_D_=6.10 across data from historical and new cohorts.

### BCG vaccination

BCG Danish Strain 1331 (Statens Serum Institut, Copenhagen, Denmark) was expanded, frozen at approximately 3 × 108 CFUs ml−1 in single-use aliquots and stored at −80 °C. Immediately before injection, BCG (for all vaccine routes) was thawed and diluted in cold PBS containing 0.05% tyloxapol (Sigma-Aldrich) and 0.002% antifoam Y-30 (Sigma-Aldrich) to prevent clumping of BCG and foaming during aerosolization. IV BCG was injected into the left saphenous vein in a volume of 2 ml; No loss of viability was observed for BCG after aerosolization. Text refers to nominal BCG doses.

### Mycobacterium tuberculosis challenge

Macaques were challenged by bronchoscope with 4–36 CFUs barcoded *Mtb* Erdman 6 months after BCG vaccination (Supplementary Table 1) in a 2 ml volume as previously described. Infectious doses across this range result in similar levels of TB disease in unvaccinated rhesus in this and previous studies. Clinical monitoring included regular monitoring of appetite, behavior and activity, weight, erythrocyte sedimentation rate, Mtb growth from gastric aspirate and coughing. These signs, as well as PET–CT characteristics, were used as criteria in determining whether a macaque met the humane end point before the predetermined study endpoint.

### Necropsy and bacterial burden

NHPs were euthanized 12 weeks after Mtb or at a humane endpoint by sodium pentobarbital injection, followed by gross examination for pathology. A published scoring system (*79*) was used to determine total pathology from each lung lobe (number and size of lesions), LN (size and extent of necrosis), and extrapulmonary compartments (number and size of lesions). All granulomas and other lung pathologies, all thoracic LNs, and peripheral LNs were matched to the final PET–CT scan and collected for quantification of Mtb. Each lesion (including granulomas, consolidations and clusters of granulomas) in the lung, all thoracic LNs, random sampling (50%) of each of the 7 lung lobes, 3–5 granulomas (if present) or random samples (30%) of spleen and liver, and any additional pathologies were processed to comprehensively quantify bacterial burdens. Suspensions were plated on 7H11 agar (Difco) and incubated at 37 °C with 5% CO2 for 3 weeks for CFU enumeration or formalin-fixed and paraffin-embedded for histological examination. CFUs were counted and summed to calculate the total thoracic bacterial burden for the macaque (*79–81*)

### Rhesus bronchoalveolar (BAL) profiling and sequencing

BAL wash fluid (3 × 20 ml washes of PBS) was centrifuged and cells were combined before counting, as previously described (*82*). High-throughput single-cell mRNA sequencing by Seq-Well was performed on single-cell suspensions obtained from NHP BAL, as previously described (*16*, *83*). Approximately 15,000 viable cells per sample were applied directly to the surface of a Seq-Well device. At each time point after BCG, two arrays were run for each sample—one unstimulated and one stimulated overnight with 20 μg/mL of PPD in R10. Sequencing for all samples was performed on an Illumina Nova-Seq.

### Data preparation

Libraries were aligned using Drop-seq tools against the M. mulatta genome using STAR50 with forced detection of 5,000 or 10,000 barcodes. Cells were then identified per array using emptyDrops (DropletUtils, R) with an FDR cutoff of 0.01. Scrublet was also utilized on each array and cells labeled as doublets or in the top 99th percentile were removed. Data from Darrah et al. was accessed on GEO (GSE139598). Digital gene expression matrices and metadata were merged and used to generate a Seurat object for all downstream analyses. We removed cells with less than 400 genes detected, less than 700 UMIs, and more than 10% or 25% reads from mitochondrial or ribosomal genes, respectively. We estimated ambient, contaminating genes using the estimateAmbience function (DropletUtils, R) and excluded genes detected in the top 99.7% percentile in at least 16 of the 32 batches within the Dose cohort samples (fig. S1E). Additional filtering was performed after preliminary clustering of unstimulated and PPD-stimulated samples separately, following the procedure outlined below. Clusters defined by multiple cell lineage markers, low library metrics, and/or high expression of ribosomal or mitochondrial genes were excluded.

### Normalization, dimensionality reduction and clustering

Single-cell analyses were performed using R and the Seurat package based on previous recommendations and benchmarking (*84*). Counts were log-normalized using NormalizeData and the top 3000 variable features were selected using FindVariableFeatures (selection.method = “vst”). PCA was run on the scaled matrix on variable features only. Selection of downstream PCs was inspected using multiple methods including the “elbow” heuristic and an intrinsic dimension estimation (maxLikGlobalDimEst, intrinsicDimension, R). Batch effects were then corrected using Harmony across both stimulation and batch variables (theta = 2, sigma = 0.1, lambda = 1). Visualization of the UMAP embedding was generated using RunUMAP across 30 dimensions. Features were visualized within the UMAP embedding using hexbin density plots (bins = 64, schex, R). Clustering was performed on the shared nearest neighbor (SNN) graph (knn = 20) using either the Walktrap algorithm (steps = 4, cluster_walktrap, igraph, R) or the Leiden algorithm (leiden_find_partition, leidenbase, R). Any clusters with less than 10 cells were automatically grouped into other clusters based on their SNN connectivity (modified GroupSingletons, Seurat, R). Leiden clustering was performed automatically by scanning resolutions between a lower bound and upper bound cluster limit then optimizing the modularity across resolutions. Resulting clusters were hierarchically clustered and reordered (BuildClusterTree, Seurat, R) based on expression of all variable features. For all cells, 30 principal components were used for Harmony and 30 Harmony-corrected components were used for clustering.

### Cell annotation

Clusters were then compared by calculating the Wilcoxon Rank Sum test statistic and AUROC for all genes (wilcoxauc, presto, R). Cells were then annotated based on expert inspection of cluster features. Select cell populations were extracted and reprocessed using the procedure and parameters previously described to further annotate cell subsets. The following groups of cell populations were further detailed using this subclustering procedure: cycling cells, B and T/NK cells, alveolar macrophages and myeloid cells, myeloid cells, and T/NK cells.

Among the T and NK cells, we identified 13 distinct cell states, including *PRF1* and *GZMB*- expressing effector *CD8*+ T cells (n = 8,088 cells), *GZMK* and *EOMES*-expressing CD8+ T cells (n = 4,354), *IFNG* and *TNF*-expressing polyfunctional T cells (n = 1,460), *TIGIT*-expressing regulatory T cells (n = 497), and *IRF4* and *CD28*-expressing helper *CD4*+ T cells (n = 6,540) (Fig. 2A and 2B and Table S5). NK cells were marked by the expression of *GNLY*, *KLRC1*, and *KLRB1*. Several transcription factors implicated in T cell activation and polarization were defining markers of cell states, including *IRF4, RBPJ, BATF, NR4A3*, and *EGR2* (*30–37*). Activated T cells were primarily defined by genes associated with the expression of *JUN* and *FOSB* along with *ATF3* and several heat shock proteins, potentially representing acutely activated T cell subsets from the BAL (*85*). *CD69*, a widely-used marker of T cell state, was most expressed in activated and polyfunctional T cells (*86*, *87*).

### Cellular abundance

Cell abundance analysis was performed using three different methods. First, we calculated Kendall’s τ statistic using the macaque quaternary group as a ranked parameter. Second, we compared fractional abundances between groups using the Mann-Whitney *U* test. Lastly, we utilized a generalized linear model to model the observed counts. Counts for each cell type were extracted across specimens and the comparison variable of interest, such as quaternary group. Paired observations of a cell type and other cell types within each sample were modeled using two interaction terms, cell type * sample type and cell type * specimen. Cell type probabilities, odds ratios, and P values were then calculated using the emmeans package in R to compare the celltype * specimen model. To visualize abundance changes, we plot the odds ratio, representing the chance of observing the cell type in one condition relative to the contrasting condition, and the Benjamini– Hochberg-adjusted P value.

### T cell signatures

Gene signature scores were calculated using AddModuleScore_UCell in R (*88*). These scores represent the collective relative ranking of genes within individual cells. Lung residency genes were derived from a blood/tissue-resolved atlas (*19*), highlighted in that publication’s figure 2C. These genes are CD69, CREM2, RGS2, SLA, NFE2L2, RGS1, LMNA, RGCC, DUSP6, and SOCS1. Module 2 genes were derived from a previously published study (*9*), consisting of 173 genes.

### Identifying gene correlates

Within a specific unstimulated or stimulated cell type, Kendall’s τ, Spearman’s ρ, and Wilcoxon Rank Sum test statistic were calculated for each gene against group, CFU, and high-dose IV BCG vs. other samples, respectively. After collating these metrics, genes were further filtered based on expression (only top two expressing cell types for each gene were considered) and consistency (all metrics were directionally consistent). This was applied to unstimulated and PPD-stimulated T cells and unstimulated and PPD-stimulated AMs/MCs.

### PPD response signatures

We performed differential expression analysis (logistic regression with study and route as latent variables) between unstimulated and PPD-stimulated cells within selected cell types (Alveolar macrophages, T cells) to define the general response to PPD. We defined a PPD-response signature by taking the top 50 genes based on log2 fold change after filtering for high-confidence cell type-specific genes. Gene signature scores were calculated using AddModuleScore_UCell in R. We then summarized the score within each batch and assessed the Pearson correlation with total CFU and difference between burden groups using non-parametric Mann-Whitney U tests. To define the number of DE genes between unstimulated and PPD-stimulated samples within each T cell subset, the same procedure described above was used but within each T cell subset.

### Gene fold-change differences across groups post-PPD stimulation

To identify genes that were differentially upregulated after PPD stimulation across groups, genes were first filtered on their expression in the cell subset of interest. Only genes most-expressed in that subset were used. Additionally, the top 200 genes based on AUROC between unstimulated and PPD-stimulated samples were used. The gene expression differences and fold changes were then calculated per sample. These metrics were then correlated with group using Kendall’s τ. For visualization, the expression difference per group was then bootstrapped (two.boot, simpleboot, R) using 10,000 replicates.

### Cell-cell interactions analysis

CellChat (v1.1.3) was used on a subset of the global dataset. Cycling cells were removed and 1000 random cells were utilized per cell type. Macaque gene symbols were converted to human symbols based on 1:1 orthologs from the Ensembl BioMart database. Cell communication probabilities were computed as described (*25*) using the human database. Interaction strengths and ligand-receptor probabilities were visualized. Only the top 3 target cell-sender cell interactions were considered for each ligand-receptor calculation. Expression score represents the average expression across all cells of genes within ligand-receptor pairs minus the average expression of randomly selected control genes within representative expression bins. Ligand-receptor pairs were selected by identifying the most associated signaling genes and extracting all genes within the respective ligand-receptor pairs. The “T cell-myeloid cell interaction” signature comprised of TNFSF8, TNFRSF8, CCL20, CCR6, LTA, TNFRSF14, CD80, CD28, CTLA4, CD40LG, CD40, TNFRSF1B, TNF, MPZL1, TNFSF14, ICAM2, ITGAL, ITGB2, CD226, CD274, NCAM1, IFNG, IFNGR1, IFNGR2. Polyfunctional T cells, activated T cells, helper CD4+ T cells, mregDCs, cDC1s, neutrophils, monocytes, alveolar macrophages, and recruited macrophages were used.

### Gene set enrichment

The top 50 genes were input to Enrichr (*89*). The MSigDB Hallmark database was used. Results were visualized in R as described in figure legends.

### Statistical analysis

All reported P values are from two-sided comparisons. Pairwise comparisons were performed in all instances using the Mann-Whitney *U* test. The Benjamini-Hochberg procedure was used to correct for multiple hypothesis testing. The Kruskal-Wallis test was used to compare sample distributions. Where applicable, all comparisons were performed by averaging cellular metrics within each batch and comparing batches, known as pseudobulking. This method was performed for the expression of specific genes and signature scores. 10,000 bootstrap replicates were used where applicable. P < 0.05 was considered significant. Adjusted P values are denoted by: *, p < 0.05; **, p < 0.01; ***, p < 0.001; ****, p < 0.0001. Statistical analyses were performed in R and RStudio.

### Supplementary Materials

Materials and Methods

Fig. S1. Macaque cohort and sequencing characteristics.

Fig. S2. Cellular annotation, features, and proportions.

Fig. S3. Myeloid features and proportions.

Fig. S4. Ligand receptor score as a function of timepoint.

Fig. S5. Cellular changes across sample groups.

Fig. S6. Lung residency and module 2 scores within T and NK cells.

Fig. S7. PPD response of T cell subsets and macrophages.

Table S1. Macaque and sample metadata. Table S2. Cellular metadata.

Table S3. Cell markers, including myeloid- and T cell-specific markers

Table S4. Cell type abundances and statistics (GLM, Kendall’s τ, and Mann-Whitney *U* tests).

Table S5. Gene correlates statistics for myeloid cells.

Table S6. Cell-cell interaction probabilities.

Table S7. Gene correlates statistics for T and NK cells.

Table S8. PPD response genes for alveolar macrophages and T cells.

## Acknowledgements

We acknowledge the outstanding work of veterinary and research technicians. We thank the Bryson and Blainey lab members for discussions and feedback.

## Author contributions

Conceptualization: JMP, EBI, JMR, RJ, AKS, BDB, SMF

Methodology: MHW, TKH, PAD, MS, JMP, EBI, JMR

Investigation: MHW, TKH, MW, PAD, MS, JMP, EBI, JMR

Data curation: JMP, SKN

Formal analysis: JMP, EBI, JMR

Visualization: JMP

Resources: MHW, TKH, PAD, MS, SKN, MR, RAS, PAD, GA, JLF, AKS

Writing - original draft: JMP, EBI, JMR, AKS, SMF, BDB

Writing - review and editing: JMP, EBI, JMR, SKN, JDB, PAD, GA, JLF, AKS, SMF, BDB

Supervision: MR, RAS, PAD, GA, JLF, AKS, SMF, BDB

Funding: MR, RAS, PAD, GA, JLF, AKS, SMF, BDB

## Data and materials availability

Publicly available experimental data and metadata is available at https://fairdomhub.org/studies/1141. RNA-sequencing data have been deposited in the Gene Expression Omnibus (GEO) under accession number GSE211191. For all data analyses, we used publicly available software. All code used for this analysis is available on Github (https://www.github.com/joshpeters/ivbcgdd).

## Supplemental Figure Captions

**Supplemental Figure 1. Macaque cohort and sequencing characteristics.** (A) CFU outcomes by vaccination route and dose. (B) Distribution of CFU outcomes. Red points indicate samples with scRNA-seq profiling. (C) CFU distribution between classified burden groups based on criteria set forth in Materials and Methods. Triangle points represent samples with scRNA-seq profiling. Color represents the route of BCG vaccine administration. (D) Correlation between total CFU and lung-specific CFU. (E) Ambient gene identification thresholds by median estimated ambient fraction and the number of flagged batches. (F) Number of UMIs counted and genes detected between filtered cells across unstimulated and PPD- stimulated samples.

**Supplemental Figure 2. Cellular annotation, features, and proportions.** (A) Clustered heatmap of scaled expression values and detection rates for the top 2 marker genes by AUROC for all annotated cell types. (B) Mean proportion of each cell type in unstimulated and PPD-stimulated samples. (C) Ratios of cell types across high-dose IV, low-dose IV, and all other conditions. (D) Difference in cell proportion ratios of T cells/alveolar macrophages (AMs) and non-AM myeloid cells to AMs across groups between week 13 and 25. P < 0.05 was considered significant. Adjusted P values are denoted by: *, p < 0.05; **, p < 0.01; ***, p < 0.001; ****, p < 0.0001.

**Supplemental Figure 3. Myeloid features**. (A) UMAP embeddings of myeloid cells. Callout UMAP embedding shows subclustering of myeloid cells. (B) Distributions of tissue-resident (gray) and recruited MoMac score (red) derived from Casanova-Acebes et al. 2021 within the recruited macrophages. (C-D) Top correlated genes across unstimulated and PPD-stimulated samples with phenotypic variables: Group, (Kendall’s τ), CFU (Spearman’s ρ), and protection (Log2FC). (C) Alveolar macrophages (D) Recruited macrophages. (E) Log2FC of top genes induced after PPD-stimulation within alveolar macrophages. (F) PPD signature score in alveolar macrophages across groups within unstimulated samples. (G) PPD signature score in alveolar macrophages across groups within unstimulated samples separated by time point.

**Supplemental Figure 4. Ligand Receptor Signaling.** Cell signaling expression score across unstimulated and PPD-stimulated samples at weeks 13 and 25.

**Supplemental Figure 5. Cellular changes across sample groups**. (A-B) Kendall’s τ correlation with Group and adjusted P values for each subset in (A) unstimulated samples (B) PPD-stimulated samples. (C) Odds ratio for each subset and Group comparison. Point shape indicates significance and color represents the contrast used.

**Supplemental Figure 6. Lung residency and module 2 scores within T and NK cells.** Lung residency scores across T and NK states within unstimulated samples. Signature is derived from Travaglini et al. 2020.

**Supplemental Figure 7. T cell and macrophage responses.** (A) Log2FC of top genes induced after PPD- stimulation. (B) PPD gene signature score across groups in stimulated samples. (C) Log2FC of PPD scores between groups across each T cell state. Adjusted P values were corrected using the Benjamini–Hochberg procedure. The Kruskal-Wallis test was used. (D) Log2FC of top genes induced after PPD-stimulation within alveolar macrophages. (E) PPD signature score in alveolar macrophages across groups within unstimulated samples. (F) PPD signature score in alveolar macrophages across groups within unstimulated samples separated by time point.

## REFERENCES

1. T. Cernuschi, S. Malvolti, E. Nickels, M. Friede, Bacillus Calmette-Guérin (BCG) vaccine: A global assessment of demand and supply balance. Vaccine. 36, 498–506 (2018).

2. World Health Organization, BCG vaccine: WHO position paper, February 2018 – Recommendations. Vaccine. 36, 3408–3410 (2018).

3. L. A. J. O’Neill, M. G. Netea, BCG-induced trained immunity: can it offer protection against COVID-19? Nat. Rev. Immunol. 20, 335–337 (2020).

4. P. Venkatesan, Progress in tuberculosis vaccine research. Lancet Microbe. 2, e12 (2021).

5. R. L. Kinsella, D. X. Zhu, G. A. Harrison, A. E. Mayer Bridwell, J. Prusa, S. M. Chavez, C. L. Stallings, Perspectives and Advances in the Understanding of Tuberculosis. Annu. Rev. Pathol. Mech. Dis. 16, 377–408 (2021).

6. P. Andersen, T. J. Scriba, Moving tuberculosis vaccines from theory to practice. Nat. Rev. Immunol. 19, 550–562 (2019).

7. C. Nunes-Alves, M. G. Booty, S. M. Carpenter, P. Jayaraman, A. C. Rothchild, S. M. Behar, In search of a new paradigm for protective immunity to TB. Nat. Rev. Microbiol. 12, 289– 299 (2014).

8. R. Kuan, K. Muskat, B. Peters, C. S. Lindestam Arlehamn, Is mapping the BCG vaccine-induced immune responses the key to improving the efficacy against tuberculosis? J. Intern. Med. 288, 651–660 (2020).

9. P. A. Darrah, J. J. Zeppa, P. Maiello, J. A. Hackney, M. H. Wadsworth, T. K. Hughes, S. Pokkali, P. A. Swanson, N. L. Grant, M. A. Rodgers, M. Kamath, C. M. Causgrove, D. J. Laddy, A. Bonavia, D. Casimiro, P. L. Lin, E. Klein, A. G. White, C. A. Scanga, A. K. Shalek, M. Roederer, J. L. Flynn, R. A. Seder, Prevention of tuberculosis in macaques after intravenous BCG immunization. Nature. 577, 95–102 (2020).

10. S. Sharpe, A. White, C. Sarfas, L. Sibley, F. Gleeson, A. McIntyre, R. Basaraba, S. Clark, G. Hall, E. Rayner, A. Williams, P. D. Marsh, M. Dennis, Alternative BCG delivery strategies improve protection against Mycobacterium tuberculosis in non-human primates: Protection associated with mycobacterial antigen-specific CD4 effector memory T-cell populations. Tuberc. Edinb. Scotl. 101, 174–190 (2016).

11. E. B. Irvine, A. O’Neil, P. A. Darrah, S. Shin, A. Choudhary, W. Li, W. Honnen, S. Mehra, D. Kaushal, H. P. Gideon, J. L. Flynn, M. Roederer, R. A. Seder, A. Pinter, S. Fortune, G. Alter, “Robust IgM responses following vaccination are associated with prevention of Mycobacterium tuberculosis infection in macaques” (2021), p. 2021.05.06.442979, doi:10.1101/2021.05.06.442979.

12. P. A. Darrah, J. J. Zeppa, C. Wang, E. B. Irvine, A. N. Bucsan, M. A. Rodgers, S. Pokkali, J. A. Hackney, M. Kamath, A. G. White, H. J. Borish, L. J. Frye, J. Tomko, K. Kracinovsky, P. L. Lin, E. Klein, C. A. Scanga, G. Alter, S. M. Fortune, D. A. Lauffenburger, J. L. Flynn, R. A. Seder, P. Maiello, M. Roederer, Airway T cells are a correlate of i.v. Bacille Calmette-Guerin-mediated protection against tuberculosis in rhesus macaques. Cell Host Microbe, S1931-3128(23)00199–3 (2023).

13. H. P. Gideon, T. K. Hughes, C. N. Tzouanas, M. H. Wadsworth, A. A. Tu, T. M. Gierahn, J. M. Peters, F. F. Hopkins, J.-R. Wei, C. Kummerlowe, N. L. Grant, K. Nargan, J. Y. Phuah, H. J. Borish, P. Maiello, A. G. White, C. G. Winchell, S. K. Nyquist, S. K. C. Ganchua, A. Myers, K. V. Patel, C. L. Ameel, C. T. Cochran, S. Ibrahim, J. A. Tomko, L. J. Frye, J. M. Rosenberg, A. Shih, M. Chao, E. Klein, C. A. Scanga, J. Ordovas-Montanes, B. Berger, J. T. Mattila, R. Madansein, J. C. Love, P. L. Lin, A. Leslie, S. M. Behar, B. Bryson, J. L. Flynn, S. M. Fortune, A. K. Shalek, Multimodal profiling of lung granulomas in macaques reveals cellular correlates of tuberculosis control. Immunity. 0 (2022), doi:10.1016/j.immuni.2022.04.004.

14. D. Pisu, L. Huang, V. Narang, M. Theriault, G. Lê-Bury, B. Lee, A. E. Lakudzala, D. T. Mzinza, D. V. Mhango, S. C. Mitini-Nkhoma, K. C. Jambo, A. Singhal, H. C. Mwandumba, D. G. Russell, Single cell analysis of M. tuberculosis phenotype and macrophage lineages in the infected lung. J. Exp. Med. 218, e20210615 (2021).

15. L. Huang, E. V. Nazarova, S. Tan, Y. Liu, D. G. Russell, Growth of Mycobacterium tuberculosis in vivo segregates with host macrophage metabolism and ontogeny. J. Exp. Med. 215, 1135–1152 (2018).

16. T. M. Gierahn, M. H. Wadsworth, T. K. Hughes, B. D. Bryson, A. Butler, R. Satija, S. Fortune, J. C. Love, A. K. Shalek, Seq-Well: portable, low-cost RNA sequencing of single cells at high throughput. Nat. Methods. 14, 395–398 (2017).

17. T. K. Hughes, M. H. Wadsworth, T. M. Gierahn, T. Do, D. Weiss, P. R. Andrade, F. Ma, B. J. de Andrade Silva, S. Shao, L. C. Tsoi, J. Ordovas-Montanes, J. E. Gudjonsson, R. L. Modlin, J. C. Love, A. K. Shalek, Second-Strand Synthesis-Based Massively Parallel scRNA-Seq Reveals Cellular States and Molecular Features of Human Inflammatory Skin Pathologies. Immunity. 53, 878–894.e7 (2020).

18. I. Korsunsky, N. Millard, J. Fan, K. Slowikowski, F. Zhang, K. Wei, Y. Baglaenko, M. Brenner, P. Loh, S. Raychaudhuri, Fast, sensitive and accurate integration of single-cell data with Harmony. Nat. Methods. 16, 1289–1296 (2019).

19. K. J. Travaglini, A. N. Nabhan, L. Penland, R. Sinha, A. Gillich, R. V. Sit, S. Chang, S. D. Conley, Y. Mori, J. Seita, G. J. Berry, J. B. Shrager, R. J. Metzger, C. S. Kuo, N. Neff, I. L. Weissman, S. R. Quake, M. A. Krasnow, A molecular cell atlas of the human lung from single-cell RNA sequencing. Nature. 587, 619–625 (2020).

20. M. Casanova-Acebes, E. Dalla, A. M. Leader, J. LeBerichel, J. Nikolic, B. M. Morales, M. Brown, C. Chang, L. Troncoso, S. T. Chen, A. Sastre-Perona, M. D. Park, A. Tabachnikova, M. Dhainaut, P. Hamon, B. Maier, C. M. Sawai, E. Agulló-Pascual, M. Schober, B. D. Brown, B. Reizis, T. Marron, E. Kenigsberg, C. Moussion, P. Benaroch, J. A. Aguirre-Ghiso, M. Merad, Tissue-resident macrophages provide a pro-tumorigenic niche to early NSCLC cells. Nature. 595, 578–584 (2021).

21. T. Hussell, T. J. Bell, Alveolar macrophages: plasticity in a tissue-specific context. Nat. Rev. Immunol. 14, 81–93 (2014).

22. R. Rajakariar, M. Hilliard, T. Lawrence, S. Trivedi, P. Colville-Nash, G. Bellingan, D. Fitzgerald, M. M. Yaqoob, D. W. Gilroy, Hematopoietic prostaglandin D2 synthase controls the onset and resolution of acute inflammation through PGD2 and 15-deoxyΔ12–14 PGJ2. Proc. Natl. Acad. Sci. 104, 20979–20984 (2007).

23. A. M. Cadena, J. L. Flynn, S. M. Fortune, The Importance of First Impressions: Early Events in Mycobacterium tuberculosis Infection Influence Outcome. mBio. 7, e00342–16.

24. A. Iwasaki, R. Medzhitov, Control of adaptive immunity by the innate immune system. Nat. Immunol. 16, 343–353 (2015).

25. S. Jin, C. F. Guerrero-Juarez, L. Zhang, I. Chang, R. Ramos, C.-H. Kuan, P. Myung, M. V. Plikus, Q. Nie, Inference and analysis of cell-cell communication using CellChat. Nat. Commun. 12, 1088 (2021).

26. J. Lai, S. Mardiana, I. G. House, K. Sek, M. A. Henderson, L. Giuffrida, A. X. Y. Chen, K. L. Todd, E. V. Petley, J. D. Chan, E. M. Carrington, A. M. Lew, B. J. Solomon, J. A. Trapani, K. Kedzierska, M. Evrard, S. J. Vervoort, J. Waithman, P. K. Darcy, P. A. Beavis, Adoptive cellular therapy with T cells expressing the dendritic cell growth factor Flt3L drives epitope spreading and antitumor immunity. Nat. Immunol. 21, 914–926 (2020).

27. M. S. Molina, J. Stokes, E. Hoffman, R. J. Simpson, E. Katsanis, J. Immunol., in press.

28. J. Ampudia, W. W. Young-Greenwald, J. Badrani, S. Gatto, A. Pavlicek, T. Doherty, S. Connelly, C. T. Ng, J. Immunol., in press.

29. A. W. Zimmerman, B. Joosten, R. Torensma, J. R. Parnes, F. N. van Leeuwen, C. G. Figdor, Long-term engagement of CD6 and ALCAM is essential for T-cell proliferation induced by dendritic cells. Blood. 107, 3212–3220 (2006).

30. R. C. Lynn, E. W. Weber, E. Sotillo, D. Gennert, P. Xu, Z. Good, H. Anbunathan, J. Lattin, R. Jones, V. Tieu, S. Nagaraja, J. Granja, C. DeBourcy, R. Majzner, A. T. Satpathy, S. R. Quake, M. Monje, H. Chang, C. L. Mackall, c-Jun overexpression in CAR T cells induces exhaustion resistance. Nature. 576, 293–300 (2019).

31. M. V. Wagle, S. J. Vervoort, M. J. Kelly, W. Van Der Byl, T. J. Peters, B. P. Martin, L. G. Martelotto, S. Nüssing, K. M. Ramsbottom, J. R. Torpy, D. Knight, S. Reading, K. Thia, L. A. Miosge, D. R. Howard, R. Gloury, S. S. Gabriel, D. T. Utzschneider, J. Oliaro, J. D. Powell, F. Luciani, J. A. Trapani, R. W. Johnstone, A. Kallies, C. C. Goodnow, I. A. Parish, Antigen-driven EGR2 expression is required for exhausted CD8+ T cell stability and maintenance. Nat. Commun. 12, 2782 (2021).

32. H. Seo, E. González-Avalos, W. Zhang, P. Ramchandani, C. Yang, C.-W. J. Lio, A. Rao, P. G. Hogan, BATF and IRF4 cooperate to counter exhaustion in tumor-infiltrating CAR T cells. Nat. Immunol. 22, 983–995 (2021).

33. M. Delacher, C. Schmidl, Y. Herzig, M. Breloer, W. Hartmann, F. Brunk, D. Kägebein, U. Träger, A.-C. Hofer, S. Bittner, D. Weichenhan, C. D. Imbusch, A. Hotz-Wagenblatt, T. Hielscher, A. Breiling, G. Federico, H.-J. Gröne, R. M. Schmid, M. Rehli, J. Abramson, M. Feuerer, Rbpj expression in regulatory T cells is critical for restraining TH2 responses. Nat. Commun. 10, 1621 (2019).

34. Y. Maekawa, Y. Minato, C. Ishifune, T. Kurihara, A. Kitamura, H. Kojima, H. Yagita, M. Sakata-Yanagimoto, T. Saito, I. Taniuchi, S. Chiba, S. Sone, K. Yasutomo, Notch2 integrates signaling by the transcription factors RBP-J and CREB1 to promote T cell cytotoxicity. Nat. Immunol. 9, 1140–1147 (2008).

35. Y. Maekawa, C. Ishifune, S. Tsukumo, K. Hozumi, H. Yagita, K. Yasutomo, Notch controls the survival of memory CD4+ T cells by regulating glucose uptake. Nat. Med. 21, 55–61 (2015).

36. H. Hosokawa, E. V. Rothenberg, How transcription factors drive choice of the T cell fate. Nat. Rev. Immunol. 21, 162–176 (2021).

37. E. Jennings, T. A. E. Elliot, N. Thawait, S. Kanabar, J. C. Yam-Puc, M. Ono, K.-M. Toellner, D. C. Wraith, G. Anderson, D. Bending, Nr4a1 and Nr4a3 Reporter Mice Are Differentially Sensitive to T Cell Receptor Signal Strength and Duration. Cell Rep. 33, 108328 (2020).

38. E. Y. Chiang, P. E. de Almeida, D. E. de Almeida Nagata, K. H. Bowles, X. Du, A. S. Chitre, K. L. Banta, Y. Kwon, B. McKenzie, S. Mittman, R. Cubas, K. R. Anderson, S. Warming, J. L. Grogan, CD96 functions as a co-stimulatory receptor to enhance CD8+ T cell activation and effector responses. Eur. J. Immunol. 50, 891–902 (2020).

39. P. P. Ahern, C. Schiering, S. Buonocore, M. J. McGeachy, D. J. Cua, K. J. Maloy, F. Powrie, Interleukin-23 Drives Intestinal Inflammation through Direct Activity on T Cells. Immunity. 33, 279–288 (2010).

40. S. A. Khader, G. K. Bell, J. E. Pearl, J. J. Fountain, J. Rangel-Moreno, G. E. Cilley, F. Shen, S. M. Eaton, S. L. Gaffen, S. L. Swain, R. M. Locksley, L. Haynes, T. D. Randall, A. M. Cooper, IL-23 and IL-17 in the establishment of protective pulmonary CD4+ T cell responses after vaccination and during Mycobacterium tuberculosis challenge. Nat. Immunol. 8, 369– 377 (2007).

41. R. Gopal, Y. Lin, N. Obermajer, S. Slight, N. Nuthalapati, M. Ahmed, P. Kalinski, S. A. Khader, IL-23-dependent IL-17 drives Th1-cell responses following Mycobacterium bovis BCG vaccination. Eur. J. Immunol. 42, 364–373 (2012).

42. G. Castro, X. Liu, K. Ngo, A. De Leon-Tabaldo, S. Zhao, R. Luna-Roman, J. Yu, T. Cao, R. Kuhn, P. Wilkinson, K. Herman, M. I. Nelen, J. Blevitt, X. Xue, A. Fourie, W.-P. Fung-Leung, RORγt and RORα signature genes in human Th17 cells. PLoS ONE. 12, e0181868 (2017).

43. The role of CD30 and CD153 (CD30L) in the anti-mycobacterial immune response - ClinicalKey, (available at https://www.clinicalkey.com/#!/content/playContent/1-s2.0-S1472979216302876?returnurl= https://%2F%2Flinkinghub.elsevier.com%2Fretrieve%2Fpii%2FS1472979216302876%3Fshowall%3Dtrue&referrer=).

44. P. K. Sharma, E. B. Wong, R. J. Napier, W. R. Bishai, T. Ndung’u, V. O. Kasprowicz, D. A. Lewinsohn, D. M. Lewinsohn, M. C. Gold, High expression of CD26 accurately identifies human bacteria-reactive MR1-restricted MAIT cells. Immunology. 145, 443–453 (2015).

45. A. Fröhlich, J. Kisielow, I. Schmitz, S. Freigang, A. T. Shamshiev, J. Weber, B. J. Marsland, A. Oxenius, M. Kopf, IL-21R on T Cells Is Critical for Sustained Functionality and Control of Chronic Viral Infection. Science. 324, 1576–1580 (2009).

46. S. S. Cheekatla, D. Tripathi, S. Venkatasubramanian, P. Paidipally, E. Welch, A. R. Tvinnereim, R. Nurieva, R. Vankayalapati, IL-21 Receptor Signaling Is Essential for Optimal CD4+ T Cell Function and Control of Mycobacterium tuberculosis Infection in Mice. J. Immunol. 199, 2815–2822 (2017).

47. B. H. Gern, K. N. Adams, C. R. Plumlee, C. R. Stoltzfus, L. Shehata, A. O. Moguche, K. Busman-Sahay, S. G. Hansen, M. K. Axthelm, L. J. Picker, J. D. Estes, K. B. Urdahl, M. Y. Gerner, TGFβ restricts expansion, survival, and function of T cells within the tuberculous granuloma. Cell Host Microbe. 29, 594–606.e6 (2021).

48. D. De Silva, L. Ferguson, G. H. Chin, B. E. Smith, R. A. Apathy, T. L. Roth, F. Blaeschke, M. Kudla, A. Marson, N. T. Ingolia, J. H. Cate, Robust T cell activation requires an eIF3- driven burst in T cell receptor translation. eLife. 10, e74272 (2021).

49. G. M. zu Horste, C. Wu, C. Wang, L. Cong, M. Pawlak, Y. Lee, W. Elyaman, S. Xiao, A. Regev, V. K. Kuchroo, RBPJ Controls Development of Pathogenic Th17 Cells by Regulating IL-23 Receptor Expression. Cell Rep. 16, 392–404 (2016).

50. A. Opejin, A. Surnov, Z. Misulovin, M. Pherson, C. Gross, C. A. Iberg, I. Fallahee, J. Bourque, D. Dorsett, D. Hawiger, A Two-Step Process of Effector Programming Governs CD4+ T Cell Fate Determination Induced by Antigenic Activation in the Steady State. Cell Rep. 33, 108424 (2020).

51. I. Comerford, W. Litchfield, E. Kara, S. R. McColl, PI3Kγ Drives Priming and Survival of Autoreactive CD4+ T Cells during Experimental Autoimmune Encephalomyelitis. PLOS ONE. 7, e45095 (2012).

52. N. Ladygina, S. Gottipati, K. Ngo, G. Castro, J.-Y. Ma, H. Banie, T. S. Rao, W.-P. Fung-Leung, PI3Kγ kinase activity is required for optimal T-cell activation and differentiation. Eur. J. Immunol. 43, 3183–3196 (2013).

53. Z. Zhao, S. Yu, D. C. Fitzgerald, M. Elbehi, B. Ciric, A. M. Rostami, G.-X. Zhang, IL-12R beta 2 promotes the development of CD4+CD25+ regulatory T cells. J. Immunol. Baltim. Md 1950. 181, 3870–3876 (2008).

54. S. J. Szabo, A. S. Dighe, U. Gubler, K. M. Murphy, Regulation of the interleukin (IL)-12R beta 2 subunit expression in developing T helper 1 (Th1) and Th2 cells. J. Exp. Med. 185, 817–824 (1997).

55. T. Bergsbaken, M. J. Bevan, P. J. Fink, Local Inflammatory Cues Regulate Differentiation and Persistence of CD8+ Tissue-Resident Memory T Cells. Cell Rep. 19, 114–124 (2017).

56. D. P. Simmons, H. N. Nguyen, E. Gomez-Rivas, Y. Jeong, A. F. Chen, J. K. Lange, G. S. Dyer, P. Blazar, B. E. Earp, A. M.P. (AMP) R. Network, D. A. Rao, E. Y. Kim, M. B. Brenner, “SLAMF7 engagement super-activates macrophages in acute and chronic inflammation” (2020), p. 2020.11.05.368647,, doi:10.1101/2020.11.05.368647.

57. J. Chen, M.-C. Zhong, H. Guo, D. Davidson, S. Mishel, Y. Lu, I. Rhee, L.-A. Pérez-Quintero, S. Zhang, M.-E. Cruz-Munoz, N. Wu, D. C. Vinh, M. Sinha, V. Calderon, C. A. Lowell, J. S. Danska, A. Veillette, SLAMF7 is critical for phagocytosis of haematopoietic tumour cells via Mac-1 integrin. Nature. 544, 493–497 (2017).

58. K. D. Mayer-Barber, B. B. Andrade, S. D. Oland, E. P. Amaral, D. L. Barber, J. Gonzales, S. C. Derrick, R. Shi, N. P. Kumar, W. Wei, X. Yuan, G. Zhang, Y. Cai, S. Babu, M. Catalfamo, A. M. Salazar, L. E. Via, C. E. Barry III, A. Sher, Host-directed therapy of tuberculosis based on interleukin-1 and type I interferon crosstalk. Nature. 511, 99–103 (2014).

59. S. Nair, J. P. Huynh, V. Lampropoulou, E. Loginicheva, E. Esaulova, A. P. Gounder, A. C. M. Boon, E. A. Schwarzkopf, T. R. Bradstreet, B. T. Edelson, M. N. Artyomov, C. L. Stallings, M. S. Diamond, Irg1 expression in myeloid cells prevents immunopathology during M. tuberculosis infection. J. Exp. Med. 215, 1035–1045 (2018).

60. E. Kaufmann, J. Sanz, J. L. Dunn, N. Khan, L. E. Mendonça, A. Pacis, F. Tzelepis, E. Pernet, A. Dumaine, J.-C. Grenier, F. Mailhot-Léonard, E. Ahmed, J. Belle, R. Besla, B. Mazer, I. L. King, A. Nijnik, C. S. Robbins, L. B. Barreiro, M. Divangahi, BCG Educates Hematopoietic Stem Cells to Generate Protective Innate Immunity against Tuberculosis. Cell. 172, 176–190.e19 (2018).

61. S. J. C. F. M. Moorlag, Y. A. Rodriguez-Rosales, J. Gillard, S. Fanucchi, K. Theunissen, B. Novakovic, C. M. de Bont, Y. Negishi, E. T. Fok, L. Kalafati, P. Verginis, V. P. Mourits, V. A. C. M. Koeken, L. C. J. de Bree, G. J. M. Pruijn, C. Fenwick, R. van Crevel, L. A. B. Joosten, I. Joosten, H. Koenen, M. M. Mhlanga, D. A. Diavatopoulos, T. Chavakis, M. G. Netea, BCG Vaccination Induces Long-Term Functional Reprogramming of Human Neutrophils. Cell Rep. 33, 108387 (2020).

62. E. Mata, R. Tarancon, C. Guerrero, E. Moreo, F. Moreau, S. Uranga, A. B. Gomez, D. Marinova, M. Domenech, F. Gonzalez-Camacho, M. Monzon, J. Badiola, J. Dominguez-Andres, J. Yuste, A. Anel, A. Peixoto, C. Martin, N. Aguilo, Pulmonary BCG induces lung-resident macrophage activation and confers long-term protection against tuberculosis. Sci. Immunol. 6, eabc2934 (2021).

63. B. Cirovic, L. C. J. de Bree, L. Groh, B. A. Blok, J. Chan, W. J. F. M. van der Velden, M. E. J. Bremmers, R. van Crevel, K. Händler, S. Picelli, J. Schulte-Schrepping, K. Klee, M. Oosting, V. A. C. M. Koeken, J. van Ingen, Y. Li, C. S. Benn, J. L. Schultze, L. A. B. Joosten, N. Curtis, M. G. Netea, A. Schlitzer, BCG Vaccination in Humans Elicits Trained Immunity via the Hematopoietic Progenitor Compartment. Cell Host Microbe. 28, 322–334.e5 (2020).

64. M. G. Netea, J. Domínguez-Andrés, L. B. Barreiro, T. Chavakis, M. Divangahi, E. Fuchs, L. A. B. Joosten, J. W. M. van der Meer, M. M. Mhlanga, W. J. M. Mulder, N. P. Riksen, A. Schlitzer, J. L. Schultze, C. Stabell Benn, J. C. Sun, R. J. Xavier, E. Latz, Defining trained immunity and its role in health and disease. Nat. Rev. Immunol. 20, 375–388 (2020).

65. R. J. W. Arts, S. J. C. F. M. Moorlag, B. Novakovic, Y. Li, S.-Y. Wang, M. Oosting, V. Kumar, R. J. Xavier, C. Wijmenga, L. A. B. Joosten, C. B. E. M. Reusken, C. S. Benn, P. Aaby, M. P. Koopmans, H. G. Stunnenberg, R. van Crevel, M. G. Netea, BCG Vaccination Protects against Experimental Viral Infection in Humans through the Induction of Cytokines Associated with Trained Immunity. Cell Host Microbe. 23, 89–100.e5 (2018).

66. M. Divangahi, P. Aaby, S. A. Khader, L. B. Barreiro, S. Bekkering, T. Chavakis, R. van Crevel, N. Curtis, A. R. DiNardo, J. Dominguez-Andres, R. Duivenvoorden, S. Fanucchi, Z. Fayad, E. Fuchs, M. Hamon, K. L. Jeffrey, N. Khan, L. A. B. Joosten, E. Kaufmann, E. Latz, G. Matarese, J. W. M. van der Meer, M. Mhlanga, S. J. C. F. M. Moorlag, W. J. M. Mulder, S. Naik, B. Novakovic, L. O’Neill, J. Ochando, K. Ozato, N. P. Riksen, R. Sauerwein, E. R. Sherwood, A. Schlitzer, J. L. Schultze, M. H. Sieweke, C. S. Benn, H. Stunnenberg, J. Sun, F. L. van de Veerdonk, S. Weis, D. L. Williams, R. Xavier, M. G. Netea, Trained immunity, tolerance, priming and differentiation: distinct immunological processes. Nat. Immunol. 22, 2–6 (2021).

67. P. Li, R. Spolski, W. Liao, L. Wang, T. L. Murphy, K. M. Murphy, W. J. Leonard, BATF-JUN is critical for IRF4-mediated transcription in T cells. Nature. 490, 543–546 (2012).

68. A. Harberts, C. Schmidt, J. Schmid, D. Reimers, F. Koch-Nolte, H.-W. Mittrücker, F. Raczkowski, Interferon regulatory factor 4 controls effector functions of CD8+ memory T cells. Proc. Natl. Acad. Sci. 118 (2021), doi:10.1073/pnas.2014553118.

69. M. C. Amezcua Vesely, P. Pallis, P. Bielecki, J. S. Low, J. Zhao, C. C. D. Harman, L. Kroehling, R. Jackson, W. Bailis, P. Licona-Limón, H. Xu, N. Iijima, P. S. Pillai, D. H. Kaplan, C. T. Weaver, Y. Kluger, M. S. Kowalczyk, A. Iwasaki, J. P. Pereira, E. Esplugues, N. Gagliani, R. A. Flavell, Effector TH17 Cells Give Rise to Long-Lived TRM Cells that Are Essential for an Immediate Response against Bacterial Infection. Cell. 178, 1176–1188.e15 (2019).

70. Z. Wang, L. He, W. Li, C. Xu, J. Zhang, D. Wang, K. Dou, R. Zhuang, B. Jin, W. Zhang, Q. Hao, K. Zhang, W. Zhang, S. Wang, Y. Gao, J. Gu, L. Shang, Z. Tan, H. Su, Y. Zhang, C. Zhang, M. Li, GDF15 induces immunosuppression via CD48 on regulatory T cells in hepatocellular carcinoma. J. Immunother. Cancer. 9, e002787 (2021).

71. N. M. Ratnam, J. M. Peterson, E. E. Talbert, K. J. Ladner, P. V. Rajasekera, C. R. Schmidt, M. E. Dillhoff, B. J. Swanson, E. Haverick, R. D. Kladney, T. M. Williams, G. W. Leone, D. J. Wang, D. C. Guttridge, NF-κB regulates GDF-15 to suppress macrophage surveillance during early tumor development. J. Clin. Invest. 127, 3796–3809 (2017).

72. L. Rochette, M. Zeller, Y. Cottin, C. Vergely, GDF15: an emerging modulator of immunity and a strategy in COVID-19 in association with iron metabolism. Trends Endocrinol. Metab. 32, 875–889 (2021).

73. F. Ma, T. K. Hughes, R. M. B. Teles, P. R. Andrade, B. J. de Andrade Silva, O. Plazyo, L. C. Tsoi, T. Do, M. H. Wadsworth, A. Oulee, M. T. Ochoa, E. N. Sarno, M. L. Iruela-Arispe, E. Klechevsky, B. Bryson, A. K. Shalek, B. R. Bloom, J. E. Gudjonsson, M. Pellegrini, R. L. Modlin, The cellular architecture of the antimicrobial response network in human leprosy granulomas. Nat. Immunol. 22, 839–850 (2021).

74. J. C. Melms, J. Biermann, H. Huang, Y. Wang, A. Nair, S. Tagore, I. Katsyv, A. F. Rendeiro, A. D. Amin, D. Schapiro, C. J. Frangieh, A. M. Luoma, A. Filliol, Y. Fang, H. Ravichandran, M. G. Clausi, G. A. Alba, M. Rogava, S. W. Chen, P. Ho, D. T. Montoro, A. E. Kornberg, A. S. Han, M. F. Bakhoum, N. Anandasabapathy, M. Suárez-Fariñas, S. F. Bakhoum, Y. Bram, A. Borczuk, X. V. Guo, J. H. Lefkowitch, C. Marboe, S. M. Lagana, A. Del Portillo, E. J. Tsai, E. Zorn, G. S. Markowitz, R. F. Schwabe, R. E. Schwartz, O. Elemento, A. Saqi, H. Hibshoosh, J. Que, B. Izar, A molecular single-cell lung atlas of lethal COVID-19. Nature. 595, 114–119 (2021).

75. N. Kim, H. K. Kim, K. Lee, Y. Hong, J. H. Cho, J. W. Choi, J.-I. Lee, Y.-L. Suh, B. M. Ku, H. H. Eum, S. Choi, Y.-L. Choi, J.-G. Joung, W.-Y. Park, H. A. Jung, J.-M. Sun, S.-H. Lee, J. S. Ahn, K. Park, M.-J. Ahn, H.-O. Lee, Single-cell RNA sequencing demonstrates the molecular and cellular reprogramming of metastatic lung adenocarcinoma. Nat. Commun. 11, 2285 (2020).

76. J. L. Delahaye, B. H. Gern, S. B. Cohen, C. R. Plumlee, S. Shafiani, M. Y. Gerner, K. B. Urdahl, BCG-induced T cells shape *Mycobacterium tuberculosis* infection before reducing the bacterial burden (2019), doi:10.1101/590554.

77. E. F. McCaffrey, M. Donato, L. Keren, Z. Chen, M. Fitzpatrick, V. Jojic, A. Delmastro, N. F. Greenwald, A. Baranski, W. Graf, M. Bosse, P. K. Ramdial, E. Forgo, D. Van Valen, M. van de Rijn, S. C. Bendall, N. Banaei, A. J. C. Steyn, P. Khatri, M. Angelo, Multiplexed imaging of human tuberculosis granulomas uncovers immunoregulatory features conserved across tissue and blood (2020), doi:10.1101/2020.06.08.140426.

78. E. Esaulova, S. Das, D. K. Singh, J. A. Choreño-Parra, A. Swain, L. Arthur, J. Rangel-Moreno, M. Ahmed, B. Singh, A. Gupta, L. A. Fernández-López, M. de la L. Garcia-Hernandez, A. Bucsan, C. Moodley, S. Mehra, E. García-Latorre, J. Zuniga, J. Atkinson, D. Kaushal, M. N. Artyomov, S. A. Khader, Tc Cell Host Microbe. 29, 165–178.e8 (2021).

79. P. Maiello, R. M. DiFazio, A. M. Cadena, M. A. Rodgers, P. L. Lin, C. A. Scanga, J. L. Flynn, Rhesus Macaques Are More Susceptible to Progressive Tuberculosis than Cynomolgus Macaques: a Quantitative Comparison. Infect. Immun. 86, e00505–17.

80. J. Phuah, E. A. Wong, H. P. Gideon, P. Maiello, M. T. Coleman, M. R. Hendricks, R. Ruden, L. R. Cirrincione, J. Chan, P. L. Lin, J. L. Flynn, Effects of B Cell Depletion on Early Mycobacterium tuberculosis Infection in Cynomolgus Macaques. Infect. Immun. 84, 1301– 1311.

81. H. P. Gideon, J. Phuah, A. J. Myers, B. D. Bryson, M. A. Rodgers, M. T. Coleman, P. Maiello, T. Rutledge, S. Marino, S. M. Fortune, D. E. Kirschner, P. L. Lin, J. L. Flynn, Variability in Tuberculosis Granuloma T Cell Responses Exists, but a Balance of Pro- and Anti-inflammatory Cytokines Is Associated with Sterilization. PLOS Pathog. 11, e1004603 (2015).

82. P. A. Darrah, R. M. DiFazio, P. Maiello, H. P. Gideon, A. J. Myers, M. A. Rodgers, J. A. Hackney, T. Lindenstrom, T. Evans, C. A. Scanga, V. Prikhodko, P. Andersen, P. L. Lin, D. Laddy, M. Roederer, R. A. Seder, J. L. Flynn, Boosting BCG with proteins or rAd5 does not enhance protection against tuberculosis in rhesus macaques. Npj Vaccines. 4, 1–13 (2019).

83. H. P. Gideon, T. K. Hughes, M. H. Wadsworth, A. A. Tu, T. M. Gierahn, J. M. Peters, F. F. Hopkins, J.-R. Wei, C. Kummerlowe, N. L. Grant, K. Nargan, J. Phuah, H. J. Borish, P. Maiello, A. G. White, C. G. Winchell, S. K. Nyquist, S. K. C. Ganchua, A. Myers, K. V. Patel, C. L. Ameel, C. T. Cochran, S. Ibrahim, J. A. Tomko, L. J. Frye, J. M. Rosenberg, A. Shih, M. Chao, C. A. Scanga, J. Ordovas-Montanes, B. Berger, J. T. Mattila, R. Madansein, J. C. Love, P. L. Lin, A. Leslie, S. M. Behar, B. Bryson, J. L. Flynn, S. M. Fortune, A. K. Shalek, Multimodal profiling of lung granulomas reveals cellular correlates of tuberculosis control (2020), doi:10.1101/2020.10.24.352492.

84. P.-L. Germain, A. Sonrel, M. D. Robinson, pipeComp, a general framework for the evaluation of computational pipelines, reveals performant single cell RNA-seq preprocessing tools. Genome Biol. 21, 227 (2020).

85. J. Jain, P. G. McCaffrey, V. E. Valge-Archer, A. Rao, Nuclear factor of activated T cells contains Fos and Jun. Nature. 356, 801–804 (1992).

86. D. Cibrián, F. Sánchez-Madrid, CD69: from activation marker to metabolic gatekeeper. Eur. J. Immunol. 47, 946–953 (2017).

87. K. Shinoda, K. Tokoyoda, A. Hanazawa, K. Hayashizaki, S. Zehentmeier, H. Hosokawa, C. Iwamura, H. Koseki, D. J. Tumes, A. Radbruch, T. Nakayama, Type II membrane protein CD69 regulates the formation of resting T-helper memory. Proc. Natl. Acad. Sci. 109, 7409– 7414 (2012).

88. M. Andreatta, S. J. Carmona, UCell: Robust and scalable single-cell gene signature scoring. Comput. Struct. Biotechnol. J. 19, 3796–3798 (2021).

89. E. Y. Chen, C. M. Tan, Y. Kou, Q. Duan, Z. Wang, G. V. Meirelles, N. R. Clark, A. Ma’ayan, Enrichr: interactive and collaborative HTML5 gene list enrichment analysis tool. BMC Bioinformatics. 14, 128 (2013).

